# hESCs-derived organoids achieve liver zonation by LSEC modulation

**DOI:** 10.1101/2024.05.05.592562

**Authors:** Yuying Zhang, Chenyan Huang, Lei Sun, Lv Zhou, Kaini Liang, Yudi Niu, Bingjie Wu, Peng Zhao, Zhiqiang Liu, Xiaolin Zhou, Peng Zhang, Jianchen Wu, Jie Na, Yanan Du

**Author notes:** Correspondence: Y.D.

## Abstract

Liver zonation, essential for diverse physiological functions, is lacking in existing organoid models, hindering their ability to recapitulate liver development and pathogenesis. Addressing this gap, we explored the feasibility of achieving zonated organoid by co-culturing human embryonic stem cells (hESCs) derived hepatocytes with hESCs derived liver sinusoidal endothelial cells (LSECs) exhibiting characteristics of either the liver lobule’s pericentral (PC) or periportal (PP) regions. Introducing zonated LSECs with variable WNT2 signaling subtly regulate hepatocyte zonation, resulting in noticeable metabolic function changes. Considering the lipid metabolism variations in PC and PP organoids, we constructed biomimetic zonated non-alcoholic fatty liver disease (NAFLD) organoids and revealed that glucagon-like peptide-1 receptor agonist (GLP-1RA) target LSECs, but not hepatocytes, indicating potential therapeutic mechanisms of GLP-1RA in NAFLD alleviation. This study highlights the crucial role of non-parenchymal cells in organoids for recapitulating niche heterogeneity, offering further insights for drug discovery and *in vitro* modeling of organ heterogeneity.

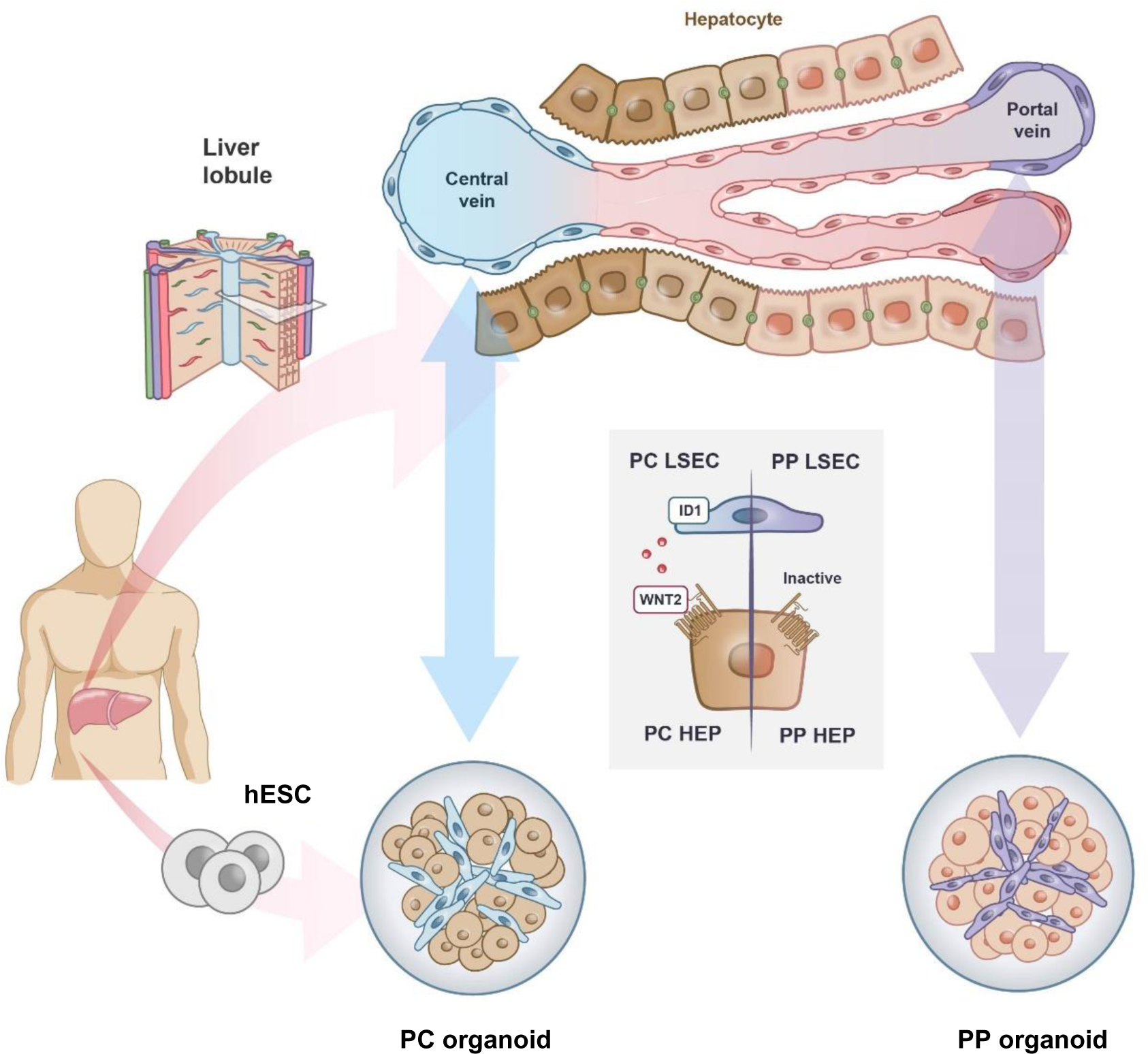
Graphical Abstract. Schematic of hESCs-derived liver zonation organoids

## Introduction

The unique architecture of liver, comprised of tens of thousands of basic units called hepatic lobules, enables it to perform numerous vital physiological functions essential for body homeostasis. The lobule is divided into periportal and pericentral areas, with mid-lobule cells in between, exhibiting spatial diversity known as zonation, characterized by variations in gene expression profiles and functional attributes among hepatocytes and non-parenchymal cells. Such non-uniform distribution of biological activities optimizes the general liver function and enables flexible response to the change of body requirements. Many liver diseases also showed zonated pathologies. For example, liver parenchymal damage resulted from drug overdose and NAFLD usually initiate from pericentral regions (Gebhardt, 1992; Yeh and Brunt, 2014), while autoimmune hepatitis and primary bile cirrhosis preferentially initiate from periportal regions (Lindor et al., 2009; Sahebjam and Vierling, 2015).

The current approach of studying the liver as a uniform entity fails to capture its intricate complexity, resulting in the loss of vital information derived from spatial heterogeneity. Even though cells in organoid models present a homogenous identity and metabolic profile, it’s imperative that these models are refined to better represent the heterogeneity of liver cells in various *in vivo* zones. Hepatocyte zonation is mainly regulated by LSEC-derived angiocrine Wnt factors (Ding et al., 2010; Goel et al., 2022; Manco and Itzkovitz, 2021). Pericentral LSECs, as the main source of angiocrine Wnt-pathway ligands (e.g., Wnt2, Wnt9b, and Rspo3), induce β-catenin activation in adjacent hepatocytes, which leads to the expression of pericentral-specific functions, such as glutamine synthesis (GS),cytochrome P450 family members Cyp2e1 (for lipid metabolism), E-cadherin (Bile acid synthesis and transport) and others (Goel et al., 2022). Specific ablation of angiocrine Wnt signaling results in decreased β-catenin activation and consequently perturbed liver metabolic zonation, as indicated by upregulation of periportal transcripts in pericentral hepatocytes (e.g., Arginase 1) (Leibing et al., 2018; Ma et al., 2020). In spite of the fact that hepatic heterogeneity has been widely investigated, it remains obscure as how LSECs, the key regulator of liver zonation, obtain their zonated profiles via various signaling cues. A better understanding of the angiodiversity of LSECs as well as the molecular progams that drive their development and zonation may pave the way for more robust differentiation of human pluripotent stem cells (hPSCs) to LSECs and, hopefully, lead to advances in success generation of liver organoids with zonated features and precision therapies for multiple liver diseases.

It can be noted that the origin of LSECs remains incompletely understood. Studies conducted in model organisms such as zebrafish, mice, birds, as well as human fetal liver samples suggest potential origins from veins, mesothelial cells, and the endodermal layer (Asahina et al., 2011; Cast et al., 2015; Crawford et al., 2010; Crivellato et al., 2007; Goldman et al., 2014; Hen et al., 2015). There are relatively few successful studies demonstrating the differentiation of LSECs from hPSCs. Koui et al. reported the first differentiation of hPSCs into cells resembling LSECs (Koui et al., 2017). They initially differentiated hPSCs into mesodermal cells, further differentiating them into endothelial progenitors. Subsequently, CD34+, CD31+, and FLK1+ LSEC progenitor cells were isolated through marker selection and were induced to overexpress LSEC markers STAB2, F8, LYVE1, and CD32B by inhibiting the TGFβ signaling pathway. Gage et al. published a second protocol for differentiating functional LSECs from hPSCs (Gage et al., 2020). They found that LSECs derived from venous endothelium exhibited higher levels of marker gene expression (e.g., CD32B and STAB2) and stronger ligand-binding capabilities. However, both studies did not compare and correlate the characteristics of hPSCs-derived LSECs with those of *in vivo* LSEC subpopulations located in different zones of the liver lobules.

A more comprehensive profiling of LSEC zonation is made possible by high-resolution spatial gene expression mapping with scRNAseq. Using paired-cell RNA sequencing that achieves spatial inference of LSECs based on the expression of landmark genes in their adjacent hepatocytes, Halpern et al. reconstructed the zonation profiles of 475 genes that are significantly zonated in LSECs (mouse) and did further validation of 12 zonated genes using smFISH (Halpern et al., 2017). Moreover, a recent study conducted by Aizarani et al. provided the single-cell transcriptome-derived zonation profiles of human LSECs, indicating limited evolutionary conservation of LSEC zonation pattern between human and mouse (Aizarani et al., 2019). In addition to well-investigated WNT2, WNT9B and RSPO3, genes such as FCN3, ICAM1 and ENG were also suggested to be part of the potential molecular signature of liver pericentral niche (Aizarani et al., 2019). Meanwhile, BTNL9 and ANPEP are thought to be potential molecular features around the periportal niche (Aizarani et al., 2019).

Acknowledging that *in vivo* compartmentalized LSECs might be one of the key regulators of hepatocyte zonation, we first generated hESCs derived PP and PC LSECs and subsequently explore the feasibility of creating 3D zonated liver organoids. 3D organoid culture system that facilitates the uniform aggregation of LSECs and hepatocytes was constructed by utilizing a polyethylene glycol (PEG) grafted polydimethylsiloxane (PDMS) microwell chip. By employing immune-fluorescence imaging and metabolic function characterization, we confirmed that co-culturing zonated LSECs with differential WNT2 signaling resulted in changes in hepatocyte zonation profiles. In an application involving zonated organoids, it was notably observed that GLP-1RA could alleviate NAFLD features by targeting GLP-1R expressed on zonated PC and PP LSECs, but not on hepatocytes. This finding provides a potential explanation for the therapeutic mechanism of GLP-1RA in the treatment of NAFLD.

## Results

### Developing efficient culture system for hESCs-derived zonated LSECs

The optimization of the method for differentiating LSECs from hESCs was based on previously published work (Gage et al., 2020; Koui et al., 2017; Zhang et al., 2021a; Zhang et al., 2021b). The basal culture medium used was a cost-effective, chemically defined homemade AATS medium, which consisted of RPMI 1640 supplemented with rice-derived recombinant human albumin, l-ascorbic acid 2-phosphate magnesium salt, human Apo-transferrin, and sodium selenite (Zhang et al., 2021a). Analysis of publicly available single-cell sequencing data (Aizarani et al., 2019) revealed that primary human PC LSECs exhibit a high expression of WNT signaling and a gene expression profile biased towards venous endothelial identity (Figure S1A and S1B), whereas primary human PP LSECs show a low expression of WNT signaling and a gene expression profile biased towards arterial endothelial identity (Su et al., 2021). Following the methodology reported by Gage et al.(Gage et al., 2020), we differentiated LSECs from hESC derived arterial and venous angioblasts separately (Figure 1A). The differentiation process was divided into three sequential stages: (1) hESCs were differentiated into mesodermal cells by the addition of BMP4 and CHIR99021 (days 0-3); (2) Subsequently, VEGF and bFGF were added to further differentiate mesodermal cells into FLK1+CD31+CD34+ LSEC progenitor cells (days 3-6); (3) Finally, the addition of TGFβ inhibitor SB431542 facilitated the maturation of LSEC progenitor cells, resulting in the expression of markers LYVE1, CD32, STAB2, and F8 in the LSECs (days 6-12) (Figure 1A). At day 6, approximately 40% of FLK1+ CD34+ CD31+ PC LSEC progenitor cells were observed in the venous-like population, whereas the arterial-like population contained around 80% of FLK1+ CD34+ CD31+ PP LSEC progenitor cells (Figure 1B). To determine the maturation of progenitor cells, we established a mCherry reporter hESC line by visualizing the expression of a common marker of the LSECs, LYVE1—using CRISPR–Cas9 genome editing. At day12, the venous-like population exhibited less than 30% differentiation of FLK1+ CD34+ CD31+ progenitor cells into LYVE1+ LSECs, whereas the efficiency in the arterial-like population approached approximately 70% (Figure 1C). Immunofluorescence staining and RNA-seq analysis revealed that both PC LSEC and PP LSEC expressed CD31, LYVE1, CD32B, STAB2, and F8 (Figure 1D and 1E). In PC EC and PC LSEC, venous marker genes (i.e., *NRP2*, *NR2F2*, *EPHB4*) showed significantly higher expression than in their PP counterparts, while arterial marker genes (i.e., *EFNB2*, *DLL4*, *NRP1*, *CXCR4*) exhibited lower expression levels (Figure 1E).In addition, we investigated features associated with LSEC zonation and observed elevated expression levels in PC LSEC for pericentral hepatocyte zonation regulators (i.e., *WNT2*, *WNT9B*, *RSPO3*), *in vivo* pericentral region high expression genes (i.e., *ICAM1*, *ENG*) and the majority of glycolysis-related metabolites (Figure 1E, 1F and S2). Besides, in PP LSEC, we observed higher expression levels of *in vivo* periportal-enriched genes, including *EFNB2* and *DLL4*, along with genes associated with oxidative phosphorylation pathways, compared to PC LSEC (Figure 1E and 1F). Moreover, immunofluorescence staining analysis also revealed significantly higher expression of EPHB4 and ICAM1 in PC EC and PC LSEC compared to their PP counterparts, while EFNB2 and CD13 exhibited lower expression levels (Fig 1G). Furthermore, the acetylated low-density lipoprotein (acLDL) uptake assay confirmed that PC LSECs phagocytose a greater amount of low-density lipid compared to PP LSECs (Figure 1H). Taken together, we validated that the PC LSECs and PP LSECs generated through VEGF pathway regulation exhibit gene expression, metabolic states, and functional differences similar to *in vivo* pericentral and periportal region LSECs.

**Figure 1.**
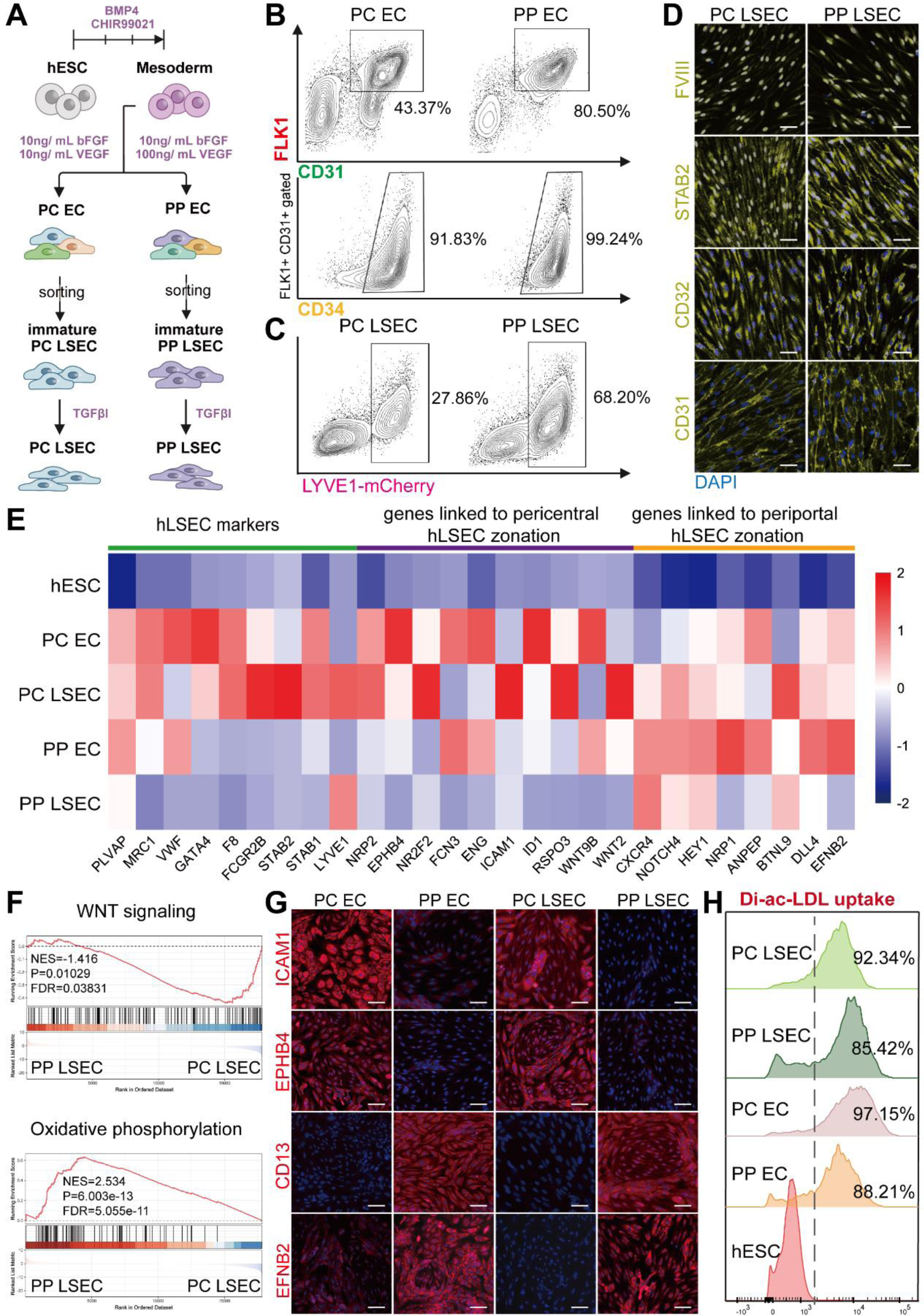
Development of efficient cell culture system for zonated hESCs-derived LSECs. (A) The protocol illustration of inducing PC and PP LSECs from hESCs; (B) The proportion of FLK1+CD31+CD34+ immature LSECs within PC and PP EC populations; (C) The proportion of LYVE1-mCherry+ LSECs within mature PC and PP LSEC populations; (D) RNA sequencing analyzed the expression of human LSEC marker genes and LSEC zonation related genes for hESC-derived PC/PP ECs and PC/PP LSECs; (E) Immunofluorescence staining of general EC marker (CD31) and three human LSEC markers (STAB2, CD32, FVIII) for hESC-derived PC/PP LSECs, with scale bar of 50 μm; (F) RNA sequencing analyzed the expression of human LSEC marker genes and LSEC zonation related genes for hESC-derived PC/PP ECs and PC/PP LSECs, PC ECs/LSECs; (G) Immunofluorescence staining of human primary PC LSEC high expressed proteins (ICAM1, EPHB4) and primary PP LSEC high expressed proteins (CD13, EFNB2) for hESC-derived PC/PP ECs and PC/PP LSECs, with scale bar of 50 μm; (H) Quantitative analysis of the proportion of cells phagocytosing DiI-Ac-LDL in hESC-derived PC/PP ECs and PC/PP LSECs.

Moreover, immunofluorescence staining analysis revealed that In PC EC and PC LSEC, EPHB4 and ICAM1 showed significantly higher expression than in their PP counterparts, while EFNB2 and CD13 exhibited lower expression levels (Fig 1G).

### 3D liver organoids formation within PEG-grafted PDMS microwell chip

To engineer and culture the liver organoids formed by hESC-derived zonated LSECs and hepatocytes, we created a low-attachment micro-well chip (Figure S3). The micro-well chip is comprised of two components: an array of two-layer wells made from PDMS (Figure 2A), which serves to confine the geometric shape of the introduced cells and matrix, and PEG modification on the surface of PDMS, enabling low adhesion and promoting the formation of spherical organoids (Figure 2B). The process of PEG graftment involves first generating radicals on the PDMS surface under UV light exposure, followed by the polymerization reaction between PEGDA monomers and PDMS (Figure 2B). From the contact angle measurements, it was evident that the surface hydrophilicity of PDMS progressively increases over a grafting time of 0 to 4 h, reaching a minimum contact angle of around 11 degrees (Figure 2C). The spherical organoids formed in the micro-well chip grafted for 4 h were uniform and regular, indicating that the PEG surface could diminish cell attachment and ECM adhesion (Figure 2D and 2E). The diameters of the spherical organoids were approximately 900 μm, accompanied by an aspect ratio of about 0.93 and a Young’s modulus of around 600 Pa (Figure 2F and 2G). By examining through scanning electron microscopy, it was observed that within liver organoids, hepatocytes and endothelial cells are layered and stacked on top of each other both on the outer and inner layers of the extracellular matrix (Figure 2H). Furthermore, immunofluorescence staining analysis revealed that CD31+ liver sinusoidal endothelial cells are able to connect with each other within the organoid spheres to form a preliminary vascular network, while ALB+ hepatocytes are uniformly distributed within the organoid spheres (Figure 2I). In summary, we contended that PEG-grafted PDMS micro-well chips exhibit excellent low-attachment propensity and biocompatibility attributes, rendering them suitable for forming and cultivating 3D liver organoids.

**Figure 2.**
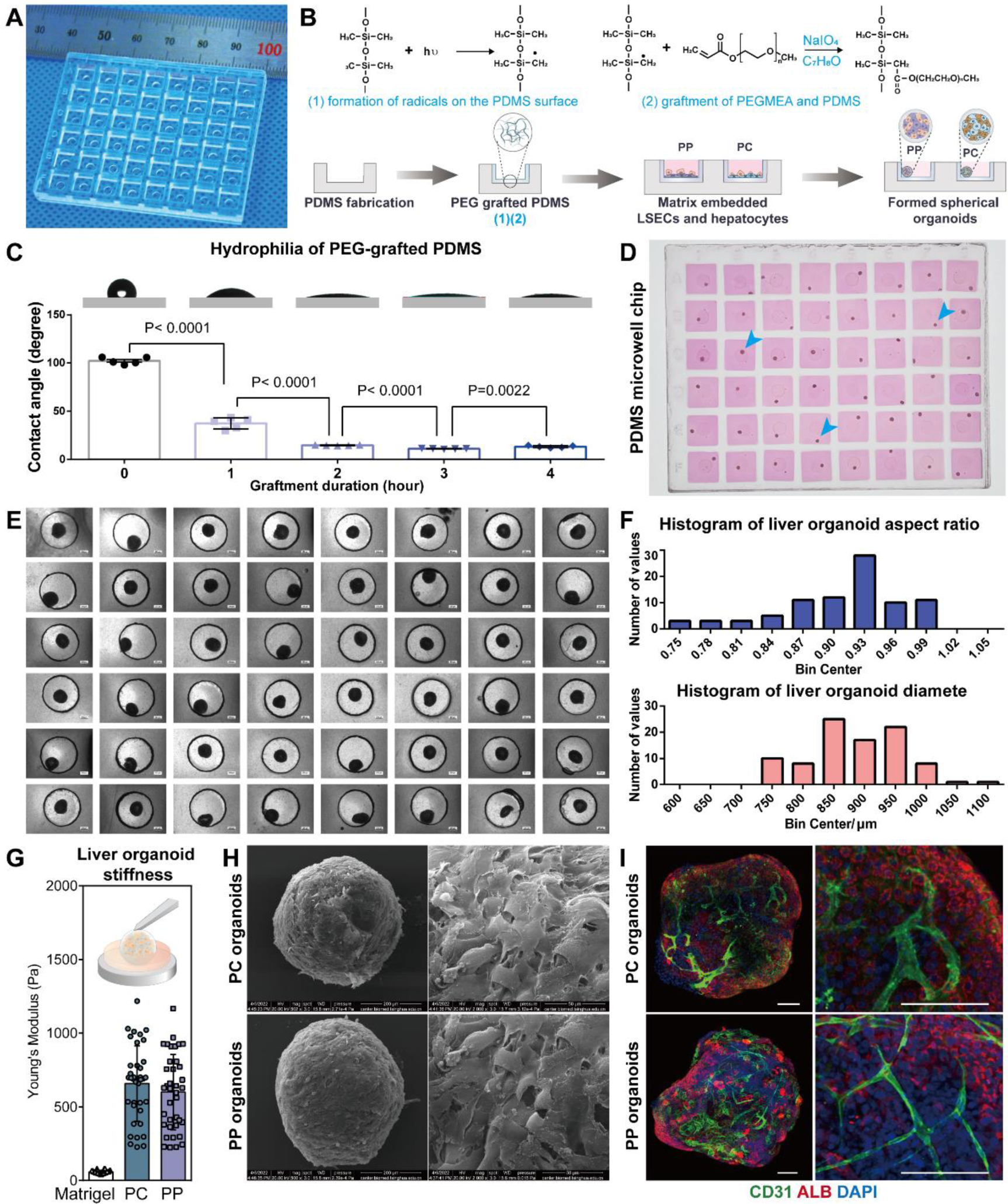
PEG-grafted PDMS microwell chips for low-attachment liver organoid culture. (A) Photo of the PDMS chip; (B) Schematic illustration of the operation process of the 3D culture system: 1) fabrication of the PDMS chip; 2) PEG grafting onto PDMS; 3) assembling hESC-derived PC/PP LSECs and hepatocytes with Collagen I and Matrigel; 4) formation of PC/PP organoids; (C) Contact angle measurement of PDMS surfaces with varying grafting times, from 0 to 4 h. Measurements were conducted in accordance with ASTM D5946-17, using a 2 μL water droplet; (D) Image of 3D organoids cultured on a PEG-grafted PDMS microwell chip, with one organoid in each well. The blue arrows indicate the organoids; (E) Enlarged bright-field images showing organoid formation 48 h after cell seeding, with a 500 μm scale bar; (F) Histogram analysis of spheroid dimensions. This histogram presents the distribution of aspect ratios and diameters for a total of ninety-two spheroids. The x-axis represents the measured aspect ratios and diameters, while the y-axis counts the number of spheroids falling within each specified range; (G) Determination of 3D PC and PP organoid stiffness by AFM; (H) SEM observation for the cell and ECM distribution of PC and PP organoids; (I) 3D visualization of immunofluorescence staining for CD31 (indicating vascular structure) and ALB (indicating hepatocyte function) in PC and PP organoids, with a scale bar of 200 μm;

### hESCs-derived organoids achieve hepatocyte zonation by WNT signaling modulation from zonated PC and PP LSECs

To investigate whether the phenotype of hepatocytes can be regulated by endothelial phenotype differences through cell-cell interactions during co-cultivation, we co-cultured hESCs-derived hepatocytes separately with hESCs-derived PC LSECs and PP LSECs on PDMS microchips to generate PC organoids and PP organoids mimicking *in vivo* liver PC and PP zone. Quantitative PCR analysis unveiled differential expression patterns in two types of organoids, showing elevated levels of downstream genes of the WNT2 signaling pathway (*LGR5*, *CTNNB1*, *AXIN2*) and *in vivo* pericentral markers (*GLUL*, *CYP2E1*, *CDH2*) in PC liver organoids compared to PP liver organoids, while *in vivo* periportal markers (*ARG1*, *CDH1*) exhibited lower expression in PC liver organoids as opposed to PP liver organoids (Figure 3A). Through immunofluorescence staining analysis of protein expression levels, we found that downstream of the WNT2 signaling pathway (β-catenin) and *in vivo* pericentral markers (GS and N-cadherin) showed higher expression in PC liver organoids compared to PP liver organoids, while in PP liver organoids, *in vivo* periportal markers (ARG1 and E-cadherin) exhibited higher expression than in PC liver organoids (Figure 3B). RNA sequencing analysis revealed that PC organoids demonstrated enhanced glutamine synthesis (e.g., *GLUL*, *GLUD1*, *GLS*), lipogenesis (e.g., *PPARA*, *FASN*, *SCD*, *ACLY*, *SREBF*, *APOA*, *APOE*), CYP450 enzyme activity (e.g., *CYP1A1*, *CYP2C9*, *CYP2C19*, *CYP3A5*), and other xenobiotic metabolism (e.g., *GSTP1*, *SULT1A1*, *NAT2*) compared to PP organoids. Conversely, PP organoids showed superior protein secretion (e.g., *SEC23 and 24*, *SEC62 and 63*, *SEC31*, *SEC22*, *ABCC1*, *SLC22*) and ureagenesis, notably in the final step of converting arginine to urea through *ARG1*, in relation to PC liver organoids. In addition, the two types of organoids reveal distinct metabolic differences in hepatocyte function with PC liver organoids exhibiting enhanced enzyme metabolism of CYP3A4 and CYP2E1 compared to PP liver organoids (Fig 3D and 3E), while PP liver organoids show a superior ability to convert ammonium chloride into urea and alcohol into aldehyde relative to PC liver organoids (Fig3F and 3G). Furthermore, the two types of organoids exhibited variations in damage response under distinct pathological conditions. Since the PC organoid exhibited higher CYP2E1 activity (Figure 3E), it converts carbon tetrachloride (CCl₄) into highly reactive trichloromethyl free radicals(·CCl3) and trichloromethyl peroxide(·CCl3O2), leading to lower cell viability and reduced albumin secretion in PC liver organoids compared to PP liver organoids (Figure 3H and 3I). While the PP organoid has higher alcohol dehydrogenase (ADH) activity (Figure 3G), converting allyl alcohol (AA) into toxic acrolein, this led to lower cell viability and reduced albumin secretion in PP liver organoids compared to PC liver organoids (Figure 3H and 3I). In summary, the regulation of liver-like organogenesis by endothelium exhibited zonal characteristics at the gene/protein expression levels, physiological functions and responses to hepatic toxins that closely resemble those of hepatic cells within the *in vivo* niche (Figure 3J).

**Figure 3.**
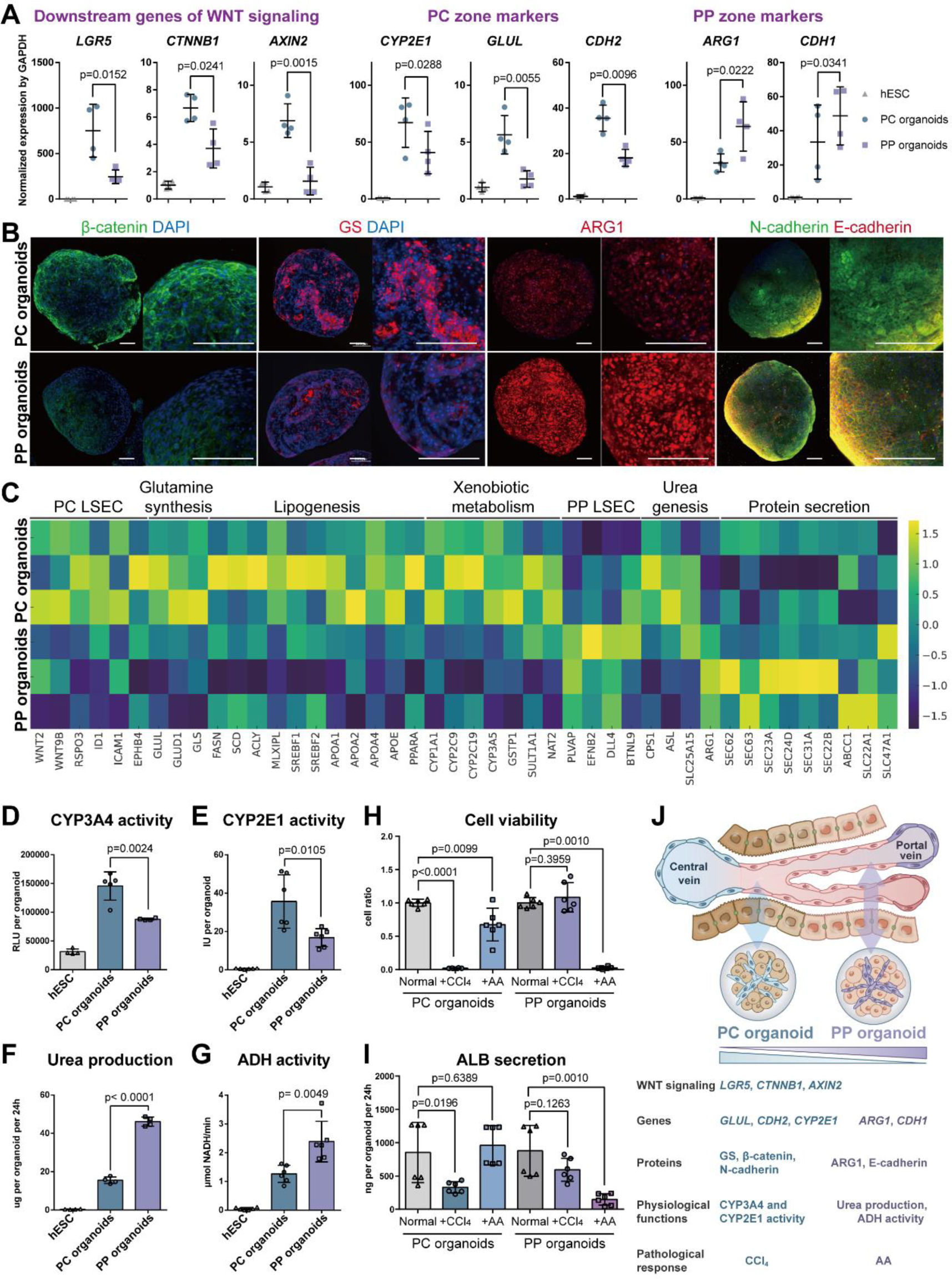
hPSCs-derived organoids achieve hepatocyte zonation. (A) qPCR results for the expression of downstream genes of the WNT/β-catenin signaling pathway (*LGR5*, *CTNNB1*, *AXIN2*), primary liver PC zone markers (*GLUL*, *CYP2E1* and *CDH2*) and primary liver PP zone markers (*ARG1* and *CDH1*) in PC and PP organoids, with *GAPDH* as a housekeeping gene. (B) Immunofluorescence staining of primary PC zone markers (β-catenin, N-cadherin) and primary PP zone markers (ARG1, E-cadherin) for PC and PP organoids, with scale bar of 200 μm; (C) RNA sequencing analyzed genes related to *in vivo* peri-central zone function (glutamine synthesis, lipogenesis, xenobiotic metabolism) and *in vivo* peri-potal zone function (urea genesis, protein secretion) of PC and PP organoids; (D) Measurement of the CYP3A4 activity for PC and PP organoids. Relative light units (RLU) per organoid is indicated; (E) Measurement of the CYP2E1 activity for PC and PP organoids. The enzyme activity is indicated in International Units (IU) per organoid. (F) Measurement of urea production in PC and PP organoids. Urea amount per organoid is indicated. (G) Measurement of ADH enzyme activity in PC and PP organoids. Enzyme activity levels that expressed as the amount of NADH (in micromoles) produced per min per organoid is indicated. (H) Cell viability assays of PC and PP organoids in medium with 4 mM carbon tetrachloride (CCl_4_) and 100 μM allyl alcohol (AA) for 24h. The absorbance ratios of each group, relative to the control group in normal medium, are indicated; (I) Albumin secretion level of PC and PP organoids in medium with 4 mM CCl_4_ and 100 μM AA for 24h. Albumin amount per organoid is indicated. (J) This schematic delineates the differences between PC and PP organoids across five key aspects: WNT signaling, marker gene expression, marker protein expression, physiological function and pathological response.

To further verify the effect of LSEC secretory signals on hepatocyte zonation, co-culture experiments of hESCs-derived LSECs and hepatocytes in a trans-well system were conducted. It was observed that co-culture with hESCs-derived PC LSECs or addition of WNT agonists CHIR99021 led to the upregulation of downstream genes (i.e., *LGR5*, *CTNNB1*, *AXIN2*) of the WNT2 signaling pathway and *in vivo* pericentral marker genes (*CYP2E1*) in hepatocytes, while co-culture with hESCs-derived PP LSECs or addition of WNT inhibitor IWP2 led to the upregulation of *in vivo* periportal marker genes (*ARG1*) (Figure 4A). Additionally, co-culturing with hESCs-derived PC LSECs in the presence of WNT inhibitor IWP2 can also activated the pericentral marker gene *CYP2E1*, suggesting that there exist other factors secreted by LSECs besides WNT signals to regulate the zonation phenotype in hepatocytes (Figure 4A). Regarding the mechanisms underlying the interplay between LSECs and hepatocytes in organoids, since ID1 has been demonstrated to mediate the upregulation of WNT2 signaling in hepatic sinusoidal endothelium during liver regeneration (Ding et al., 2010), we hypothesize that hESC-derived LSECs may potentially regulate the zonal phenotype of hepatic cells akin to the *in vivo* scenario by modulating the secretion of WNT2 signal through the transcription factor ID1. Both gene and protein expression levels of ID1 and WNT2 signaling were upregulated in PC LSECs compared to PP LSECs (Figure 4B and 4C). To investigate the impact of altered ID1 expression levels on WNT2 expression, we generated conditions of ID1 downregulation using an inhibitor (AGX51) and facilitated ID1 overexpression using a doxycycline-inducible system in hESCs. Western blot analysis demonstrated that reducing ID1 expression in hESC-derived endothelial cells by AGX51 resulted in decreased WNT2 expression (Figure 4D and S4A-C), while inducing ID1 overexpression in hESC-derived endothelial cells led to heightened WNT2 expression (Figure 4E and S4D-E). To assess whether low ID1 expression affects the endothelial regulation of hepatocytes, AGX51 was added to the transwell-based co-culture of hESC-derived PC LSECs with hepatocytes. This led to decreased expression of *LGR5* and increased expression of *in vivo* periportal marker gene *ARG1*, which indicates that lower ID1 expression may cause the ablation of angiocrine WNT signaling, resulting in decreased β-catenin activation and subsequently perturbed liver metabolic zonation (Figure 4A). To assess whether ID1 overexpression influences endothelial regulation of hepatocyte, PC organoids were generated from ID1 KI hESC-derived PC LSECs with and WT hESC-derived hepatocytes. Following 5 days of doxycycline induction, there was a slight increase in the expression of β-catenin in PC organoids with ID1 KI LSECs, while there was no obvious change in the PP organoids (Figure 4F). Compared to ID1 KI LSECs-formed PC organoids without DOX, those treated with DOX showed significantly increased expression of *CTNNB1*, *LGR5*, *GLUL*, *AXIN2*, *CDH2*, and slight elevations in *CYP2E1* (Figure 4G). In summary, within our engineered organoid model, hESCs-differentiated endothelial cells exert regulatory control over the hepatocyte zonation phenotype primarily through the ID1-WNT2 axis, among other mechanisms (Figure4H).

**Figure 4.**
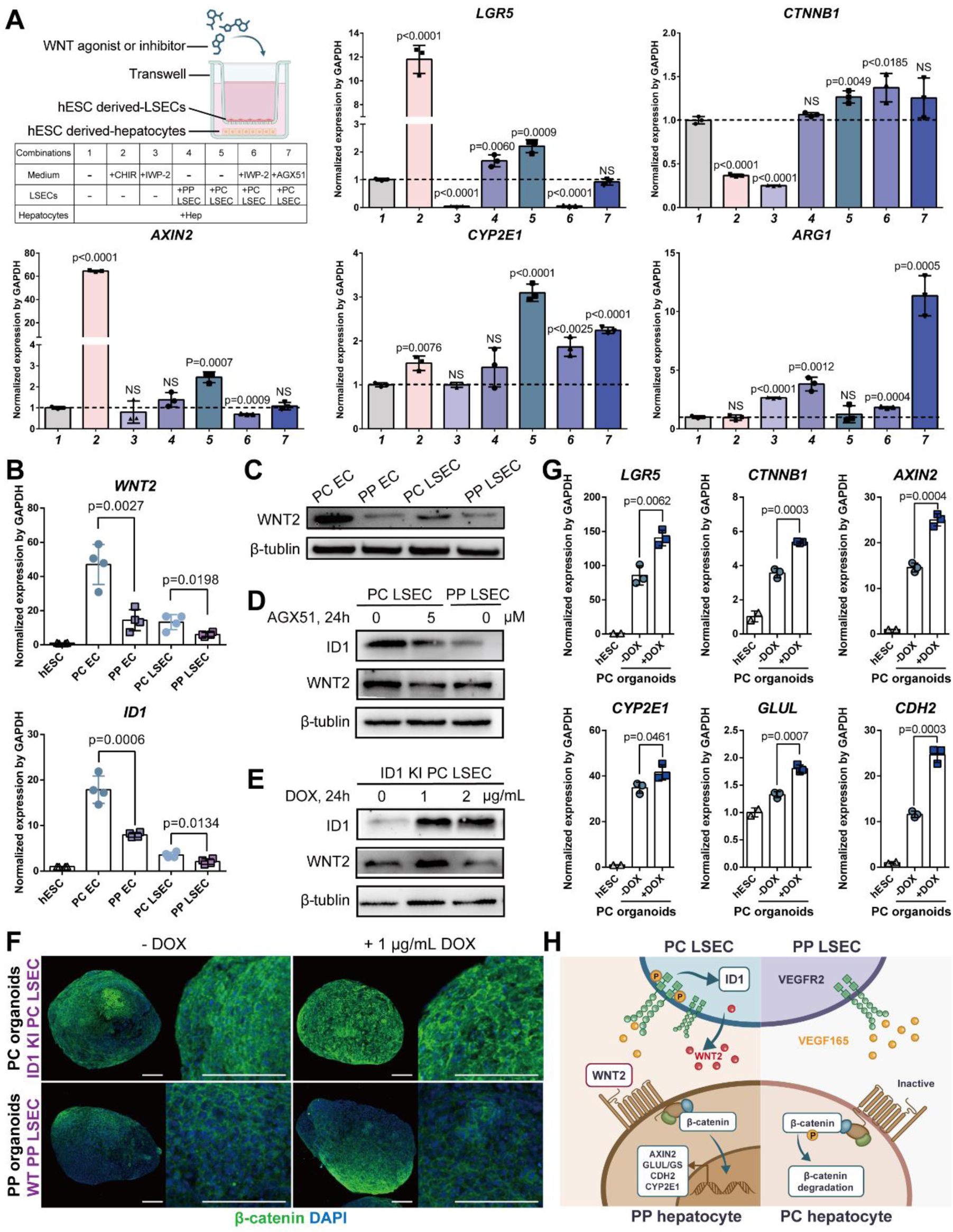
Hepatocyte zonation was modulated by WNT2 signaling from zonated PC and PP LSECs. (A) qPCR results for the expression of WNT/β-catenin signaling pathway (*LGR5*, *CTNNB1*, *AXIN2*), primary liver PC zone marker (*CYP2E1*) and primary liver PP zone marker (*ARG1*) in hESCs-derived hepatocytes that are co-cultured with hESCs-derived PC or PP LSECs and small molecular in transwell, with GAPDH as the housekeeping gene. The culture medium was supplemented with three different small molecules: CHIR99021 at 2 μM, IWP-2 at 5 μM, and AGX51 at 5 μM; (B) qPCR results for the expression of *WNT2* and *ID1* genes in hESCs-derived PC EC, PP EC, PC LSEC and PP LSEC, with *GAPDH* as the housekeeping gene; (C) Western blot analysis for WNT2 protein in hESCs-derived PC and PP LSEC, with β-tublin as the housekeeping protein. (D) Western blot analysis for ID1 and WNT2 protein in hESCs-derived PC LSEC, PP LSEC and PC LSEC treated with 5 μM ID1 inhibitor AGX51 for 24h, with β-tublin as the housekeeping protein. (E) Western blot analysis for ID1 and WNT2 protein in ID1 knock in (KI) hESCs-derived PC LSEC treated with 0, 1, and 2 μg/mL doxycycline (DOX) for 24 h, with β-tublin as the housekeeping protein. (F) Immunofluorescence staining of β-catenin for hESCs-derived PC and PP organoids, with and without treatment with 1 μg/mL DOX, with scale bar of 200 μm. PC organoids are composed of ID1 KI PC LSECs and WT hepatocytes, whereas PP organoids are composed of WT PP LSEC and WT hepatocytes, with scale bar of 200 μm; (G) qPCR results for the expression of downstream genes of the WNT/β-catenin signaling pathway (*LGR5*, *CTNNB1*, *AXIN2*) and primary liver PC zone markers (*GLUL*, *CYP2E1* and *CDH2*) in hESCs-derived PC organoids, with and without treatment with 1 μg/mL DOX, with GAPDH as the housekeeping gene. PC organoids are composed of ID1 KI PC LSECs and WT hepatocytes. (H) The schematic illustrated the interaction between LSECs and hepatocytes in PC and PP organoids. ID1-WNT2-β-catenin axis may be one of the potential underlying mechanisms by which LSECs regulate hepatocyte zonation *in vitro*.

### Establishment and application of zonated liver NAFLD organoids

Owing to the regulatory influence of WNT on liver lipid metabolism (Hall et al., 2017; Yao et al., 2018), notable differences in lipid metabolism were observed between PC organoids and PP organoids. Metabolic pathways, including fatty acid metabolism, biosynthesis of unsaturated fatty acids, PPAR signaling, cholesterol metabolism, CYP450 metabolism, glycerophospholipid metabolism and NAFLD metabolism, demonstrated increased activity in PC organoids compared to PP organoids (Figure 5A and S5A). NAFLD is a widespread disease without specific drug treatments. Multiple cell-based and animal-based (e.g., mouse model) experiments, as well as clinical trials, have suggested that GLP-1R could be a potential target for NAFLD treatment, although its mechanism of action remains unclear (Ding et al., 2010). We propose that human zonated NAFLD organoids could be utilized to verify the effects of GLP-1RA drugs and conduct related mechanistic investigation. Initially, NAFLD organoids were induced by supplementing the organoid medium with free fatty acids (FFA), consisting of 450 μM oleic acid (OA) and 150 μM palmitic acid (PA) (Figure S5B). Subsequent observations indicated that PC organoids accumulated lipid droplets at a faster rate compared to PP organoids, and the total amount of lipid droplets accumulated was also greater in PC organoids than in PP (Figure 5B). The addition of 200 μM Exendin-4 (Ex-4), a GLP-1R agonist, into both PC and PP organoids resulted in a modest reduction in the amount of lipid droplets, yet there was a notable decrease in the total amount of triglycerides (TG) contained within them (Figure 5C). It is hypothesized that GLP-1RA may predominantly enhance the oxidation of fatty acids and may also increase the export of TGs from the liver. Next, we developed a GLP-1R knockdown (KD) ESC cell line using lentivirus-mediated shRNA to further investigate the mechanism of GLP-1RA treatment. Western blot and quantitative PCR results indicated the presence of GLP-1R in wild-type (WT) hESCs-derived PC and PP LSECs, and the absence of GLP-1R in both WT hESCs-derived hepatocytes and GLP-1R KD hESCs-derived PC and PP LSECs (Figure 5D and 5E). Addition of Ex-4 to the culture medium activated downstream cAMP signaling (Tomas et al., 2020). Under 2D conditions, WT hESCs-derived PC LSECs produced more cAMP in response than PP LSECs, while both WT hESCs-derived hepatocytes and GLP-1R KD hESCs-derived PC and PP LSECs showed no response to the agonist (Figure S5C). Meanwhile, under NAFLD conditions induced by FFA, Ex-4 could reduce apoptosis of WT ESC-derived PC LSECs and PP LSECs, and had no effect on GLP-1R KD hESCs-derived PC and PP LSECs (Figure S5D). Furthermore, 3D organoid cultures were utilized using four LSEC subtypes, WT PC, GLP-1R KD PC, WT PP, and GLP-1R KD PP, each co-cultured with WT hepatocytes (Figure 5F). In WT PC and PP LSEC organoids, treatment of Ex-4 was observed to reduce cellular apoptosis as demonstrated by cell viability assays, improve liver function as evidenced by increased ALB secretion, reduce inflammation through decreased IL-6 production, and enhance the survival of endothelial cells as revealed by increased CD31 fluorescence signals (Figure 5G). In contrast, in GLP-1R KD PC and PP LSEC organoids, Ex-4 did not demonstrate a significant effect (Figure 5G). These comparisons confirmed that GLP-1RA targets GLP-1R expressed on LSECs, rather than directly impacting hepatocytes. In conclusion, within the liver, GLP-1RA is likely to alleviate NAFLD indirectly through activation of GLP-1R on the LSECs, rather than directly affecting hepatocytes (Figure 5H). Zonated organoids thus provide a valuable model for exploring the varied phenotypes of NAFLD and assessing the impacts and mechanism of action of various drugs.

**Figure 5.**
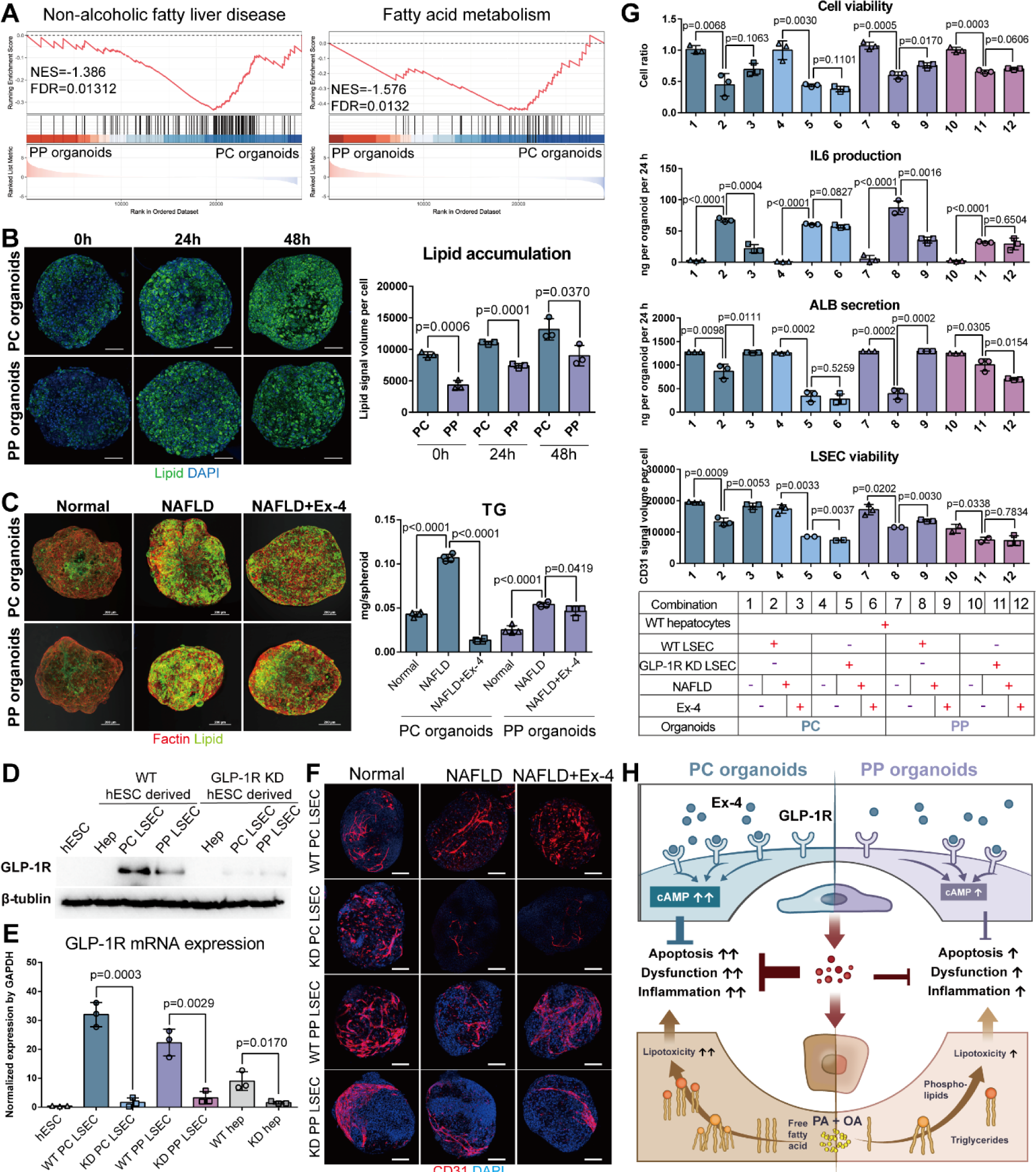
Exendin-4 alleviates cell apoptosis and inflammation in biomimetic liver zonated PC and PP NAFLD organoids through GLP-1R expressed on zonated PC and PP LSECs. (A) Gene Set Enrichment Analysis (GSEA) of NAFLD and lipid metabolism pathways in PC and PP Organoids; (B) Characterization of lipid accumulation in PC and PP organoids treated with 450 μM OA and 150 μM PA at 0, 24, and 48 h. Lipid accumulation is visualized as green fluorescence signal, quantified by dividing the volume of the green signal by the blue DAPI signal, which stains nuclei. (C) Characterization of lipid accumulation and TG level in PC and PP organoids after treatment with normal culture medium for 5 days, NAFLD (450 μM OA and 150 μM PA) culture medium for 5 days, and NAFLD culture medium supplemented with 200 nM Exendin-4 (Ex-4) for 5 days. Lipid accumulation is visualized in green, while F-actin, characterizing the cytoskeleton, is depicted in red. (D) Western blot analysis for GLP-1R protein in hESC, WT hESCs-derived PC LSEC, PP LSEC, hepatocyte (Hep), and GLP-1R KD hESCs-derived PC LSEC, PP LSEC, Hep, with β-tublin as the housekeeping protein. (E) qPCR analysis for GLP-1R protein in hESCs, WT hESCs-derived PC LSEC, PP LSEC, hepatocyte (Hep), and GLP-1R KD hESCs-derived PC LSEC, PP LSEC, Hep, with GAPDH as the housekeeping gene; (F) Immunofluorescence staining of CD31 in hESCs-derived PC and PP organoids treated with Normal, NAFLD, NAFLD + 200 nM Ex-4 medium for 5 days. PC organoids are composed of PC LSEC or GLP-1R KD PC LSECs with WT hepatocytes, whereas PP organoids are composed of PP LSEC or GLP-1R KD PP LSECs with WT hepatocytes, with scale bar of 200 μm; (G) Cell viability assays, IL6 production, albumin secretion and LSEC cell viability of hESCs-derived PC and PP organoids treated with Normal, NAFLD, NAFLD + 200 nM Ex-4 medium for 5 days. PC organoids are composed of PC LSEC or GLP-1R KD PC LSECs with WT hepatocytes, whereas PP organoids are composed of PP LSEC or GLP-1R KD PP LSECs with WT hepatocytes. For cell viability assays, the absorbance ratios of each group, relative to the corresponding control group in normal medium, are indicated. Albumin and IL6 amount per organoid are indicated. LSEC cell viability is visualized as a red CD31 fluorescence signal (Figure 5F), quantified by dividing the volume of the red signal by the blue DAPI signal, which stains nuclei. (H) The schematic illustrated the potential explanation for the therapeutic mechanism of GLP-1RA in the treatment of NAFLD PC and PP organoids.

## Discussion

Our work represents the first study to develop hPSC-derived LSECs with differential WNT2 expression, aligning with zonation-specific characteristics, and to modulate the emergence of zonal metabolic traits in hepatocytes. This work highlights the crucial role of endothelial cells in regulating liver organoid heterogeneity and mediating the mechanisms of action of GLP-1RA.

The hESC-derived LSECs we differentiated, whether directed towards arterial or venous phenotypes, were able to highly express the marker proteins of *in vivo* human LSECs. This was demonstrated not only through immunofluorescence staining experiments but also using an endogenous LYVE1-mCherry reporter system. In addition, our protocol, which takes 13 days to yield LSECs, is cost-effective, with clearly defined components, making it suitable for applications in cell therapy, large-scale organoid production, and other clinical needs. Moreover, when comparing the *in vitro* differentiated LSECs with *in vivo* human LSECs, large vessel endothelium, and other endothelial cells, we found that the venous-like LSECs most closely resemble the *in vivo* PC zone LSECs (Figure S1C). However, the currently differentiated PP LSECs are not sufficiently similar to their *in vivo* counterparts (Figure S1C). It may be necessary to explore factors beyond VEGF that influence endothelial regulation, such as the extracellular matrix, shear stress or hydrostatic pressure.

In the cultivation of 3D organoids, commercially available low-adhesion plates are convenient to use, but they have disadvantages such as being non-reusable, having unstable low-adhesion effects, being expensive, and not customizable to laboratory needs. Hinman et al. reported that the PEG-graftment methodology could effectively achieve low adsorption of large molecules such as collagen, proteins, and DNA on PDMS. (Hinman et al., 2021). We adopted this method to build PEG grafted PDMS microwell chip for low-attachment culture of cells and organoids. It should be noted that this modification method might lead to cytotoxicity, thus requiring repeated rinsing of the chip with alcohol and deionized water. This PEG grafted PDMS microwell chip demonstrates superior capabilities in promoting low cellular adhesion, preserving cell viability, and enabling the formation of organoids. Moreover, its modification allows for repeated usage, withstanding 7-10 cleaning cycles without compromising its functionality. In terms of plate design, there is a groove that can accommodate 3 μ l of liquid, making the loading of cell suspensions into wells more precise and ensuring a more uniform cell distribution. The organoids produced are uniform and controllable, suitable for large-scale operations. Overall, our design of low-adhesion plates can provide good reference for laboratories to prepare personalized plates and for the commercial large-scale production of organoid plates.

In the construction of zonated liver organoids, the basis for achieving zonated liver organoids *in vitro* depends on the understanding of *in vivo* liver cell zonation mechanisms. Currently, liver zonation mechanisms mainly include endothelial WNT signaling, the Renin-Angiotensin System (RAS) signaling (Peng et al., 2021) and glucagon (Cheng et al., 2018). In our preliminary tests, glucagon could not induce hepatocyte zonation (Figure S6A), and EGF in the RAS signaling could induce zonation of the *CDH1* gene but not the important *ARG1* gene (Figure S6A). In our study, endothelial WNT signaling induced hepatocyte zonation more effectively than the two above-mentioned methods. Moreover, although WNT proteins are secretory, their solubility is poor. Due to the absence of specific small molecule agonists for WNT2, WNT9B and RSPO3, it is currently impossible to directly add small molecules to regulate hepatocyte zonation. Our experiments show that it is possible to induce local zonation characteristics in hepatocytes by adding broad-spectrum WNT agonists or inhibitors, but the effect is no better than co-culturing with endothelial cells. For instance, the WNT agonist CHIR99021 increases CYP2E1 gene expression but does not enhance GLUL gene expression (Figure S6A); the WNT inhibitor IWP2 elevates ARG1 expression but fails to decrease CYP2E1 gene expression (Figure 4A). Beyond providing signals like WNT2, the significance of endothelial cells in organoids far exceeds that of small molecules, as evidenced by GLP-1RA studies in NAFLD organoids. Notably, GLP-1R is mainly found on endothelial cells, with minimum expression on hepatocytes, highlighting the critical role of endothelial cells in drug response evaluation. At last, to successfully develop zonated liver organoids *in vitro*, it is crucial to emphasize that the differentiation state of hepatocytes is fundamental in determining the emergence of zonation characteristics. Immature hepatocytes in the AFP+ALB+CYP3A4-ALT-stage are more easily induced to exhibit zonation characteristics, whereas it is difficult to induce zonation characteristics in mature hepatocytes with different endothelia (Figure S3B, S3C and S6A). For the liver zonation by LSEC modulation, literature reports that endothelium can also regulate the zonation of two additional liver cells: stellate cells and macrophages (Manco and Itzkovitz, 2021). In summary, to construct zonated liver organoids *in vitro* that mimic *in vivo* heterogeneity, the role of endothelium cannot be ignored, the differentiation state of various cells is crucial, and more *in vivo* zonation mechanisms need to be studied.

Based on the NAFLD model constructed using zonated liver organoids, it has been proposed that GLP-1RA drugs may exert their effects in the liver by acting on GLP-1R located on LSECs. Although the liver is considered to barely express GLP-1R, researchers have observed through HE staining that GLP-1R is expressed on liver sinusoidal endothelial cells, and its expression increases in NAFLD (Yokomori and Ando, 2020). In our investigations, GLP-1R expression was identified in isolated mouse primary LSECs (Figure S6B). This finding underpins the hypothesis that GLP-1RA could exert a direct effect on the liver, thereby suggesting a mechanism for their action beyond the traditional glucoregulatory pathways. This discovery contributes to the evolving understanding of GLP-1RA’s role in hepatic pathology and potentially in the development of GLP-1R targeted therapy for liver diseases. In addition, experiments also show that addition of Ex-4 to NAFLD liver organoids mainly reduces cell apoptosis, enhances hepatocyte function, and decreases inflammation, but does not significantly clear lipid droplets. It was observed that reducing the concentration of FFA in the culture medium proved more effective than the addition of Ex-4 in reducing lipid droplet accumulation within hepatocytes following NAFLD induction (Figure S6C). This implies that the treatment of NAFLD needs to reduce lipid in the blood, improve the immune environment of the liver, and rescue liver cell functions. For the treatment of many diseases, such as weight rebound, gastrointestinal adverse reactions, thyroid cancer, suicidal depression tendencies etc., GLP-1R is an excellent drug target, but its side effects may raise concerns. This may be due to the wide variety of cells in the body that express GLP-1R. We speculate that targeting specific diseases in specific regions for treatment, and precise medication might not only cure the disease but also reduce the occurrence of adverse reactions. In the liver, where the GLP-1R is mainly expressed on LSECs, in addition to targeting the islets and other means to reduce blood lipid levels for the treatment of NAFLD, GLP-1RA drugs also need target LSECs to improve the immune environment of the liver and rescue liver cell functions.

## Acknowledgements

We acknowledge colleagues from the School of Medicine, Tsinghua University: Prof. Jie Na for assisting with hESCs culture and differentiation into endothelial cells, as well as Huizhen Cao and Jinyu Wang in the SLSTU-Nikon Biological Imaging Center for assisting with microscope operation and image processing. **Funding:** This work was financially supported by the National Natural Science Foundation of China (82125018) and Natural Science Foundation of Beijing, China (Z230016).

## Author contributions

Y.D. and Y.Z. conceived and designed the experiments; Y.Z. performed experiments and data collection; C.H. assisted in exploring the ID1/WNT2 mechanism; L.S. assisted in WB experiments; Y.N. and X.Z. assisted in bioinformatics data analysis; K.L. assisted in schematics design and production; P.Z. assisted in liver organoids AFM test; Y.Z. and Y.D. wrote the manuscript; Y.D is the principal investigators of the supporting grants.

## Declaration of interests

The authors declare no competing interests.

## STAR★Methods

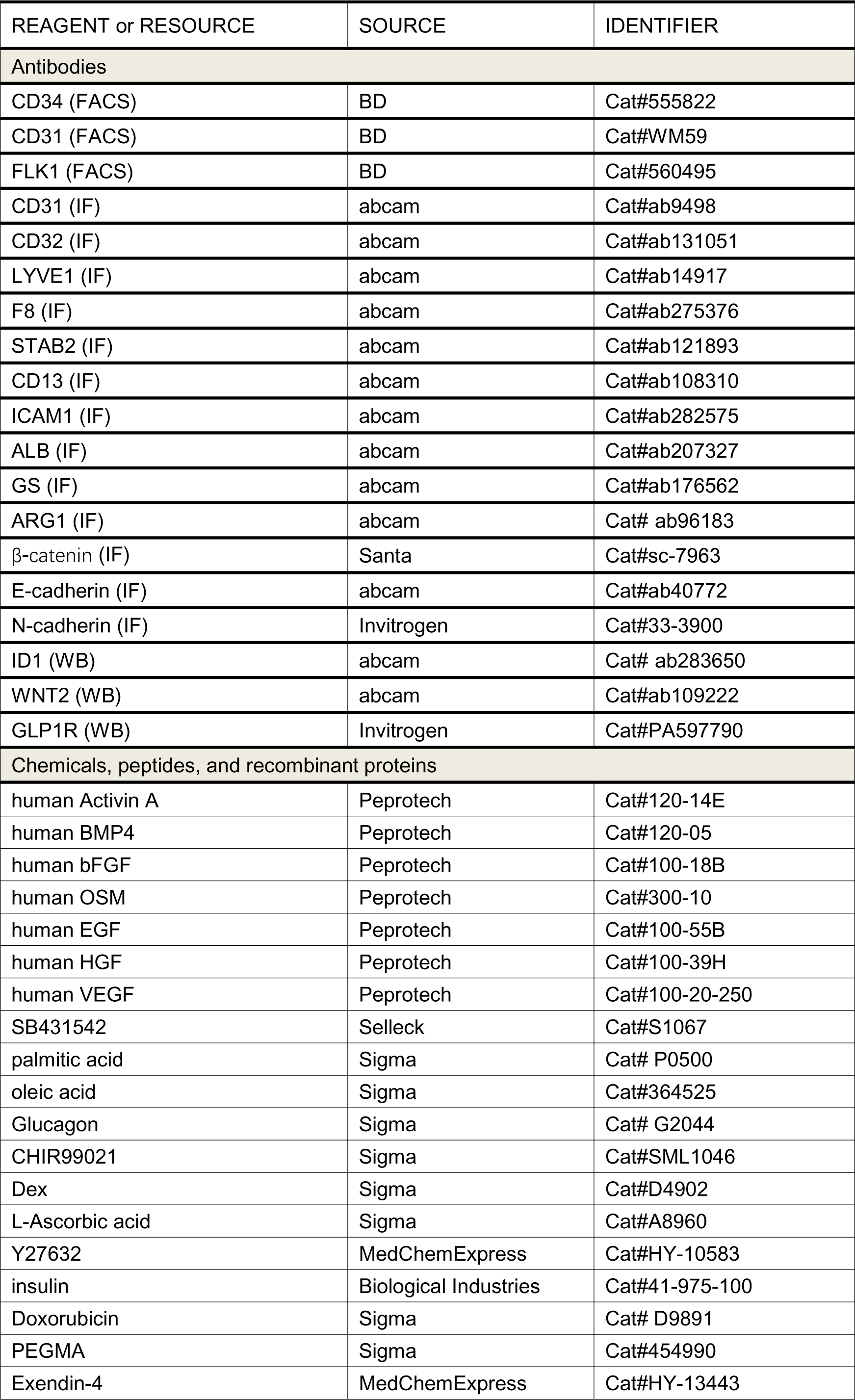
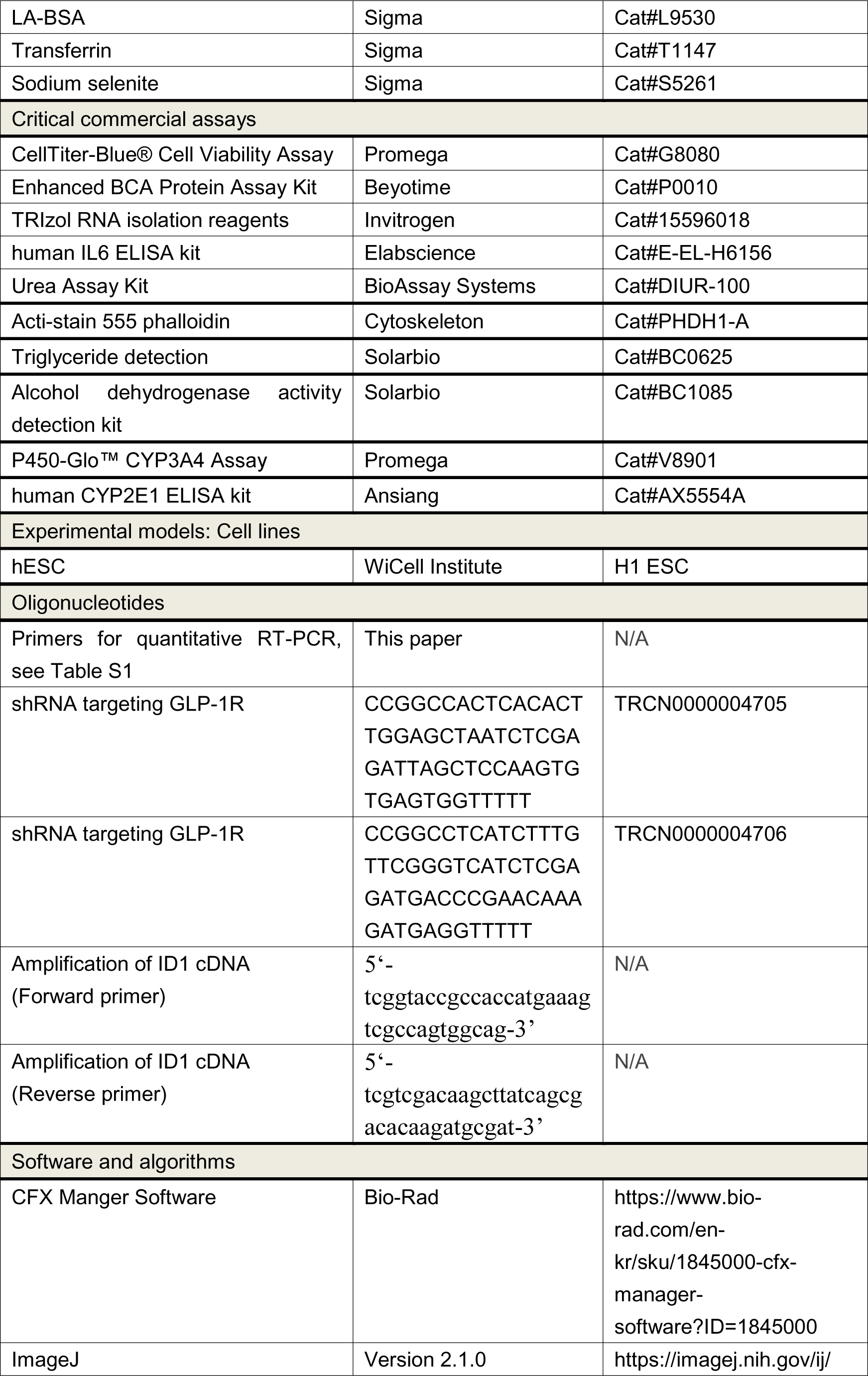
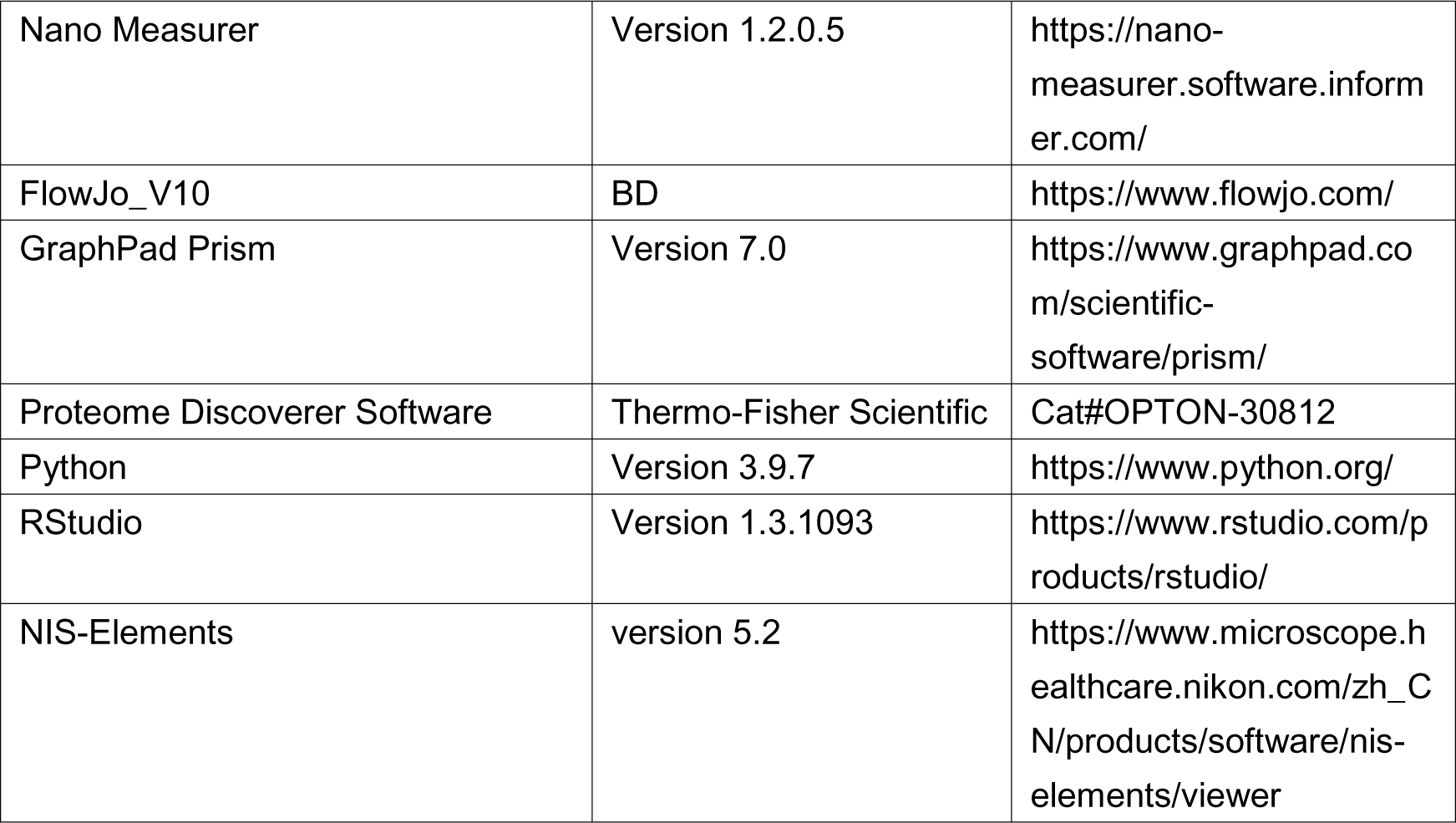
Key resources table.

Key resources table

## Experimental model and subject details

### Cell culture and differentiation

The h1 hESC lines were maintained in mTeSR1 essential medium in a 37°C incubator at 5% CO2 and 100% humidity and the medium was changed daily. hESC lines and hESC-derived cells were seeded onto Matrigel-coated plates. For initiation of cell differentiation, hESCs were seeded at a density of 1.2 × 10^5 cells per well of a 12-well plate in mTeSR1 essential medium containing 10 μM Y27632.

hESC-derived endothelial cells induction was optimized based on the method described by Zhang et al.(Zhang et al., 2021b), employing AATS medium from Professor Na Jie’s Laboratory, Tsinghua University. Initially, hESCs were differentiated into mesoderm cells by culturing in AATS medium supplemented with 5 ng/ml BMP4 for one day, followed by an additional two days with both 5 ng/ml BMP4 and 2 μM CHIR99021. For the differentiation into PC EC progenitors, mesoderm cells were dissociated using Accutase, seeded onto Matrigel-coated plates at a density of 2.3×10^5 cells per well, and cultured in AATS medium enriched with 10 ng/ml VEGF, 20 ng/ml bFGF, and B27 supplement for three days. In a parallel process, PP EC progenitors were obtained by seeding similarly treated mesoderm cells at a density of 2.8×10^5 cells per well and cultivating them in AATS medium with 100 ng/ml VEGF, 20 ng/ml bFGF, and B27 minus insulin supplement for three days. The maturation of both PC and PP progenitors into LSECs was accomplished by treating the cultures with 5 μM SB431542 for six days.

Differentiation of embryonic stem cells into Hepatocytes is divided into four stages. The process began with definitive endoderm differentiation, where hESCs were initially cultured in RPMI1640 medium with 100 ng/ml Activin A, 3 μM CHIR99021, 1% GlutaMax, and B27 supplement on Day 1. This was followed by a continuation in RPMI1640 containing 100 ng/ml Activin A, 5 ng/ml bFGF, and 1% GlutaMax, along with B27 supplement for the next two days. From Days 4 to 7, the cells were cultured in SFD medium comprising 75% IMDM, 25% Ham’s F12, 0.5% N2 supplement, 1% B27 supplement, 0.05% BSA, and 50 mg/ml ascorbic acid, supplemented with 100 ng/ml Activin A and 5 ng/ml bFGF. The subsequent stage of hepatic endoderm formation, from Days 8 to 13, involved culturing the cells in DMEM with 1 ng/mL glucose, 1% B27 supplement, 1% BSA, 50 mg/ml ascorbic acid, 40 ng/ml bFGF, and 50 ng/ml BMP4. Differentiation into hepatoblasts occurred over Days 14 to 19 in DMEM with 1 mg/mL glucose, 1% B27 supplement, 1% BSA, 50 mg/ml ascorbic acid, 20 ng/ml OSM, 25 ng/ml HGF, and 1 μM Dexamethasone. The final maturation into premature hepatocytes was carried out from Days 20 to 31 in DMEM with 5 mg/mL glucose, supplemented with 1% B27 supplement, 1% BSA, 50 mg/ml ascorbic acid, 20 ng/ml OSM, 25 ng/ml HGF, and 1 μM Dexamethasone.

### Fabrication of PEG-grafted PDMS microwell chip

The mold for the chip was fabricated using PDMS as per the design outlined in Supplementary Figure 3. PDMS was prepared by mixing the base and the curing agent in a 10:1 ratio, poured into the mold, and then degassed in a vacuum until all bubbles were eliminated. The mold was then placed in a 65°C oven for overnight curing. After curing, the PDMS was carefully demolded in 35% alcohol and washed with deionized water overnight. For low adsorption modification of the well plate surface, the method referenced Hinman et al. (Hinman et al., 2021) and involved ultraviolet (UV) cross-linking of Poly ethylene glycol (PEG). This was achieved by adding a PEG crosslinker solution containing 10% weight percentage PEG monomer, 0.5% weight percentage benzyl alcohol, and 0.5 mM sodium periodate to PDMS well chips. UV cross-linking was performed using a UV lamp, pausing every 30 min for degassing and replenishment of the solution. Post-crosslinking, the mold was thoroughly rinsed with deionized water to remove any unreacted PEG, followed by continuous washing on a shaker for three days, with daily water changes. Subsequently, the modified gel plates were sterilized in 75% alcohol for 24 h and then exposed to UV light for 3 h to ensure complete evaporation of the alcohol, rendering the plates ready for use.

### 3D liver organoids formation

To construct 3D liver organoids in vitro, day 25 hepatocytes along with PC or PP LSECs derived from hESCs were enzymatically digested into single cells. These cells were subsequently resuspended in an organoid formation matrix at a 1:1 ratio. The matrix was composed of Matrigel (3.35 mg/ml) and type I collagen (1.11 mg/ml), blended in organoid growth medium. This growth medium was a mixture of endothelial cell differentiation medium and hepatocyte medium, containing equal parts AATS medium and DMEM/F12 medium. It was further supplemented with 50 ng/ml VEGF, 20 ng/ml bFGF, 40 ng/ml HGF, 20 ng/ml EGF, 1% GlutaMax, 1% B27, 1% BSA, and 1% penicillin-streptomycin (PS). The organoid cell suspension was seeded onto a 48-well PDMS chip at a density of 30,000 cells per well, with each well receiving 3 μl of the suspension. The chip was incubated at 37 °C for approximately 45 min to facilitate matrix gelation. After gelation, organoid growth medium supplemented with 10μM Y27632 was added to each well, and the medium was refreshed daily. The cultured organoids were maintained at 37 °C in a controlled environment of 5% CO2 and 95% air. Organoids, after 5 days of formation, are suitable for a diverse range of analyses, including immunofluorescence, quantitative PCR, atomic force microscopy, scanning electron microscopy, RNA sequencing, and other techniques.

### Construction of LYVE1-mCherry reporter hESC cell line

The Cas9(sgRNA) plasmid used was pSpCas9(BB)-2A-GFP (Addgene plasmid ID: 48138), and the repair template plasmid was pL552 (Addgene plasmid ID: 68407), following the methodology outlined by Ran et al (Ran et al., 2013). The CRISPR Design Tool (http://tools.genome-engineering.org) was employed to design sgRNAs targeting the C-terminus of the LYVE1 gene, which were inserted into the PX458 plasmid. The target sequences for designed sgRNAs were 5′-gaggctggtttctttcatgc-3′ and 5′-gaaatgaggagacacacctg-3′. The homology-directed repair (HDR) template was constructed by amplifying homologous arm fragments from h1ESC’s genomic DNA, extending 1 kb from each side of the stop codon. These two homologous arm fragments were fused with mCherry, LoxP-PGK-Hyg-LoxP through high-fidelity enzyme-mediated overlap extension PCR and inserted into the pL552 plasmid. H1 ESCs were transfected with these two plasmids using Lipofectamine 3000. GFP-positive cells, indicative of successful transfection, were isolated via flow cytometry 48 h post-transfection. These cells were then cloned and expanded in a culture medium containing 50 μg/mL hygromycin (Hyg) for two weeks. PCR and flow cytometry assessments were conducted to confirm successful gene editing. PCR primers are 5‘-agagtaggtctcaaggccat-3’ (upstream) and 5‘-ctctcagtcccttcttgttgg-3’ (downstream).

Following confirmation, the cells were further transfected with pCAG-Cre: GFP (Addgene plasmid ID:13776) to facilitate recombination and removal of the Hyg segment. The final LYVE1-mCherry reporter hESC cell line was cultured in mTeSR™ medium.

### Construction of ID1 overexpression hESC cell line

To establish a Dox-controlled ID1 overexpression system in hESCs at the AAVS1 (PPP1R12C) site, the first step involved extracting total RNA from endothelial cells differentiated from hESCs. This RNA was reverse transcribed to obtain total cDNA, which served as a template. Using primers 5‘-tcggtaccgccaccatgaaagtcgccagtggcag-3’ (forward) and 5‘-tcgtcgacaagcttatcagcgacacaagatgcgat-3’ (reverse), the DNA encoding fragment corresponding to ID1 transcript variant 1 was amplified. This fragment was integrated into the pAAVS1-Neo-CAG-M2erTA backbone (Addgene plasmid ID:85798), which contains a G418 resistance gene and Tet-on system, through a double enzyme digestion and ligation process (KpnⅠ and SalⅠ), forming the donor plasmid targeted to the AAVS1 site. The CRISPR/Cas9 plasmid used in the experiment was a commercial plasmid (Addgene plasmid ID:118414) obtained by inserting the sgRNA (5‘-gTGGGGGTTAGACCCAATATC-3’) into the pX458 backbone. H1 ESCs were transfected with these two plasmids using Lipofectamine 3000 and selected with 50 μg/μl G418 commenced 72 h post-transfection and continued for two weeks. Single clones were picked from the wells, expanded in 48-well plates, and subjected to genomic PCR and sequencing for identification of correctly inserted positive clones, using primers 5‘-tcgtcgactcagcgacacaagatgcgat-3’ (forward) and 5‘-ctctggctccatcgtaagca-3’ (reverse).

### Construction of GLP-1R knockdown expression hESC cell line

The specific shRNA targeting GLP-1R was sourced from the MISSION shRNA Library. This shRNA was then cloned into the pLV-CMV-puro lentivirus mammalian expression vector, preparing it for effective delivery into hESCs. Lentivirus production was initiated by cotransfecting 293FT cells with this shRNA-laden expression vector and the second-generation lentivirus packaging vectors, pVSVG and pΔ8.9. Upon harvesting the lentiviral suspension, it was introduced into h1ESCs seeded in a six-well plate, utilizing polybrene (8 μg/ml) to enhance viral infection efficiency. To isolate and enrich for cells that successfully integrated the shRNA construct, we added puromycin at a concentration of 2.5 μg/ml to the culture medium, 24 h post-infection.

### Induction of NAFLD organoids

After 5 days of formation, organoids can also be utilized to induce NAFLD organoids, which were induced by supplementing the organoid medium with FFA. For the preparation of FFA solutions, palmitic acid (PA) and oleic acid (OA) were processed using immediate-use protocols. PA solution was prepared by dissolving 0.4 g of NaOH in 100 mL of deionized water to create a 0.1 M NaOH solution, into which 0.0256 g of PA was added. This mixture was then heated in a 75°C water bath, yielding a 40 mM PA solution. Concurrently, 1.2 g of BSA was dissolved in 2.5 mL of water and centrifuged at 8000 rpm for 5 min without shaking to form a 40% BSA solution, which was preheated to 55°C. The PA and BSA solutions were mixed in a 1:3 ratio, resulting in a 10 mM PA stock solution, which was then filtered through a 0.22 μm filter.

OA stock solution was similarly prepared by adding 31.7 μL of OA to 2ml of 0.1 M NaOH, obtaining a 50 mM OA solution. This was followed by the dissolution of 1.8 g of BSA in 5 mL of water and centrifugation at 8000 rpm for 5 min to achieve a 30% BSA solution. A 1:1 mixture of the OA and BSA solutions produced a 25 mM OA stock solution, which was also subjected to 0.22 μm filtration. Both lipid stock solutions were stored at 4°C.

## Method details

### RNA isolation and reverse transcription-PCR

RNA was extracted from both 2D cells and 3D organoids using TRIzol Reagent (Invitrogen). For reverse transcription, 500 ng of the extracted RNA was processed using the Hiscript II qRT SuperMix Kit (V) (Vazyme), converting RNA into complementary DNA (cDNA). Quantitative analysis of gene expression was performed via real-time PCR, utilizing AceQ qPCR SYBR Green Master Mix (Vazyme) on a CFX96 Real-Time System (Bio-Rad). The relative expression levels of target genes were calculated employing the 2^ΔΔCt method, with normalization against the housekeeping gene GAPDH to account for variations in cDNA input across samples. Detailed sequences of the primers used for amplification are provided in Table S1.

### Immunofluorescence staining

Cells and organoids were fixed with 4% paraformaldehyde (PFA) for 15 min, ensuring structural preservation, and then washed three times with phosphate-buffered saline (PBS) to remove any residual fixative. For permeabilization, a 0.5% Triton X-100 solution in PBS was applied for 20 min, facilitating antibody access to intracellular targets. Subsequently, cells were blocked with 5% bovine serum albumin (BSA) in PBS for 1 h to minimize nonspecific antibody binding. Overnight incubation at 4 °C with primary antibodies, previously diluted in blocking solution, was followed by several washes to remove unbound antibodies. Cells were then incubated with suitable secondary antibodies, prepared as per the primary antibodies’ specifications. Cell nuclei were stained with Hoechst 33342 for 20 min. Images were captured with a Nikon A1 HD25 confocal microscope. The fluorescence intensity of organoids and 2D cells was analyzed using NIS-Elements software version 5.2.

### Flow cytometry

To obtain suspended cells, hESC-derived PC or PP EC and LSECs were digested with accutase at 37°C for 5 min. Following digestion, the cells were suspended and incubated with primary antibodies at 37°C for 30 min. The cells were then centrifuged, the supernatant discarded, and the pellet resuspended in fresh culture medium. Subsequently, cell sorting was performed using a BD Influx sorter, and analysis was carried out with a BD Biosciences LSR Fortessa Cell Analyzer. Compensation was adjusted using single-color controls. Data analysis and result plotting were conducted using FlowJo software.

### mRNA Sequencing and Analysis

Total RNA from 2D cells and liver organoids was extracted using TRIzol and sent to Anoroad Gene Technology Corporation for sequencing. Quality was checked with a KaiaoK5500® Spectrophotometer and RNA Nano 6000 Assay Kit on a Bioanalyzer 2100. Sequencing libraries were made using the NEBNext® Ultra™ RNA Library Prep Kit for Illumina®, following the provided instructions. Data cleaning was performed with Trimmomatic to remove low-quality reads and adaptors. HISAT2 was used for aligning clean reads to the human genome. Gene counts were done with HT-seq and normalized as RPKM using Ensembl annotations and the edgeR package. DESeq2 identified differentially expressed genes. Functional enrichment analyses for GO and KEGG were conducted with the GOstats package. GSEA used the pre-ranked module in GSEA 4.03 software. mRNA sequencing for PC EC, PP EC, PC LSEC, PP LSEC and PC and PP organoids was conducted in three independent replicates. Following sequencing, comprehensive gene clustering and Gene Set Enrichment Analysis (GSEA) were performed to identify distinct gene expression pattern.

### AcLDL uptake

hESCs-derived PC or PP EC and LSECs were seeded at a density of 2.5 × 10^5 cells per well in 12-well plates. Cells were incubated with 10 µg/mL of Dil-labeled acLDL (1,1’-dioctadecyl-3,3,3’,3’-tetramethylindocarbocyanine perchlorate-labeled acetylated LDL) for 4 h at 37°C in a 5% CO2 atmosphere. Post-incubation, cells were washed thrice with PBS to remove unbound acLDL. Cellular uptake of Dil-acLDL was quantified by measuring fluorescence intensity using a BD Biosciences LSR Fortessa Cell Analyzer set at an excitation wavelength of 514 nm and an emission wavelength of 565 nm.

### Glucose metabolism analysis

To analyze glucose metabolism, we employed a non-targeted LC/MS approach following metabolite extraction from cultured cells. Initially, cells were washed with PBS and lysed in cold 80% methanol at –80°C for an hour, ensuring thorough metabolite recovery. After scraping and collecting the cell lysate, it was centrifuged to separate the supernatant, which was then use N2 to dry at room temperature and stored at –80°C until analysis.

### Contact angle measurement

Contact angle measurement was performed according to ASTM D5946-17 despite that a droplet of 2 μL was suspended. During measurement, a humidifier was used to stabilize relative humidity at 40% and the ambient temperature was 21℃. Flat PDMS samples with different grafting time were tested using the sessile drop method on OCA15Pro (Dataphysics, Germany). Deionized water was suspended on the sample and images of the water droplet was captured after the droplet was stable. Images were processed by SCA20 (Dataphysics, Germany) and Yong-Laplace fitting was used in all images to measure the contact angle. Each sample was tested by at least 4 droplets.

### Organoid Characterization

For scanning electron microscopy (SEM) analysis, liver spheroids were fixed with 2.5% glutaraldehyde in PBS at room temperature for 2 h and subsequently washed three times with PBS. The samples underwent dehydration through treatment with ethanol solutions at increasing concentrations (70%, 80%, 90%, and then 100% twice), each stage lasting for 10 min. After the removal of excess ethanol, the samples were covered with tert-butanol and stored at 4°C. They were then freeze-dried using the Tousimis Samdri-795 and coated with a 30 nm gold layer using a Quorum Q150T ES sputter coater. Electron microscopy was conducted using a Quanta 200 at an acceleration voltage of 5 keV and a working distance of 5 mm.

Atomic force microscopy (AFM) was utilized to measure the Young’s modulus, adhering to the same protocol detailed in our previously published work(Zhang et al., 2021b).

### Liver-Specific Function Assay

Following exposure to 2 mM ammonium chloride for 24 h, urea levels in organoid culture media were quantified by employing the Urea Assay Kit.

24 h after replacing the culture medium with fresh medium, collect the supernatant to measure albumin using an Albumin Detection Kit (enzyme-linked immunosorbent assay) and to measure IL-6 using an IL-6 ELISA Kit.

After replacing with fresh culture medium, CYP3A4 activity was tested by P450-Glo™ CYP3A4 Assay Kit.

After lysing the organoids by sonication in the culture medium, the supernatant was collected for CYP2E1 detection using a CYP2E1 ELISA Kit.

After the organoids are lysed by sonication with the reagents provided by the kit, the supernatant is collected to indirectly quantify ADH activity by monitoring the production of NADH using the ADH detection Kit.

After the organoids are lysed by sonication with Reagent I, provided by the kit, triglycerides in the sample undergo hydrolysis. The resultant dye reagents are then oxidized, yielding colored compounds. This process employs the Triglyceride Assay Kit.

All of procedures were executed in strict adherence to the manufacturer’s instructions.

### Cell viability evaluations

Cell viability was assessed by ATP quantifications using the CellTiter-Glo Luminescent Cell Viability Assay. Luminescence signals were measured using PerkinElmer EnSight Multimode Plate Reader.

### Analysis of publicly available single-cell sequencing datasets

Diffusion pseudo-time (dpt) analysis (Haghverdi et al., 2016) was performed, and diffusion maps were generated utilizing the destiny R package. The parameter for the number of nearest neighbors, denoted as k, was configured to 100. Subsequently, Self-Organizing Maps (SOM) were created using the FateID package, relying on the computed ordering by dpt as the input. The SOM analysis exclusively considered genes with counts exceeding 2 after size normalization in at least one cell. To outline smooth zonation profiles, a local regression was applied to normalized transcript counts following the ordering of cells by dpt. Following this, a one-dimensional SOM comprising 200 nodes was computed based on these profiles after z-transformation. In cases where the Pearson’s correlation coefficient of the average profiles for neighboring nodes surpassed 0.85, these nodes were merged. The resultant aggregated nodes constitute the gene modules illustrated in the SOM Figures.

We employed the plot-expression function in the FateID package to plot pseudo-temporal expression profiles for genes that are representative in different regions. The normalized expression data was filtered to only include the genes with the minimum expression of 2 in at least one cell. The pseudo-time ordering for endothelial cells was obtained using the same approach depicted above by retrieving the eigenvectors of the diffusion map. Outliers were removed to show the trend of gene expressions.

### Pearson correlation between bulk RNA-seq from hESCs derived ECs and scRNA-seq from human liver samples

Both unprocessed count matrices underwent cleaning and normalization procedures utilizing the RaceID package within the R programming environment. Rigorous quality control measures were implemented, involving the exclusion of cells exhibiting a comparatively low total transcript count. Specifically, for human liver cells, those with fewer than 1000 transcripts and less than 2 counts of a gene in at least one cell were omitted. Additionally, cells expressing more than 2% of KCNQ1OT1 transcripts, a previously identified marker indicative of low-quality cells, were excluded. Transcripts exhibiting a Pearson’s correlation coefficient greater than 0.4 with KCNQ1OT1 were also eliminated. In the case of bulk RNA-seq samples, cells with less than 100 transcripts and less than 1 transcript count of a gene in at least one cell were excluded due to the limited sample size, consisting of only four cells. Following the application of the aforementioned procedures, the pseudo-time analysis for Liver Sinusoidal Endothelial Cells (LSECs) was ordered, subsequently categorizing them into periportal (PP LSEC) and central (PC LSEC) clusters. Post subsetting the two datasets by shared genes, Pearson correlation coefficients were computed for each sample in bulk RNA-seq and each cluster in human liver cells, allowing for an exploration of the association between our bulk sequencing results and the standard liver cell sequencing conducted by Aizarani et al. (Aizarani et al., 2019).

## Quantification and statistical analysis

Statistical analyses were performed using one-way analysis of variance (ANOVA) with the GraphPad Prism 7.0. For experiments comparing two groups, we performed a two-tailed unpaired Student’s t test. Statistical differences were represented by p value, Data were expressed as means ± s.d. Exact P values are marked in the figures. Details of sample sizes and statistical tests for experiments are specified in the figure legends.

---

**Sup Figure 1.**
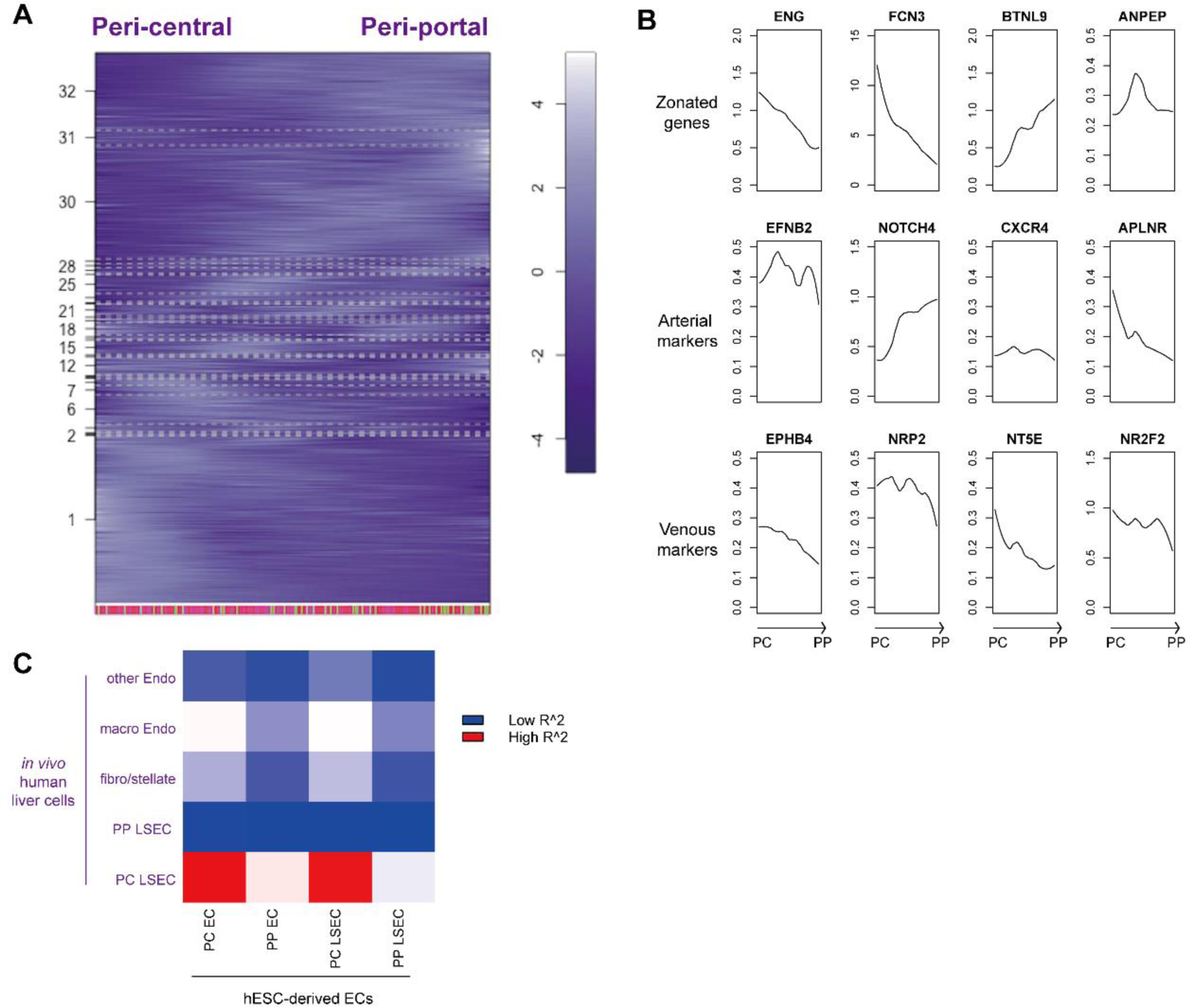
Comparative gene expression profiling of primary human PC LSECs and PP LSECs reveals differential endothelial identity, related to Figure 1. (A) Self-Organizing Maps of single-cell transcriptome-derived zonation profiles for endothelial cells (n = 1,361 cells). Colour bars at the right of SOMs show RaceID3 cluster. The gene module pattern is similar to reference paper. (B) Zonation profiles of LSEC zonation related genes (ENG, FCN3, BTNL9, ANPEP), arterial markers (EFNB2, NOTCH4, CXCR4, APLNR), and venous markers (EPHB4, NRP2, NT5E, NR2F2). The y axis of the zonation profiles indicates normalized expression. (C) Heatmap of Pearson correlation between bulk RNA-seq form hESCs derived ECs and scRNA-seq form human liver samples.

**Sup Figure 2.**
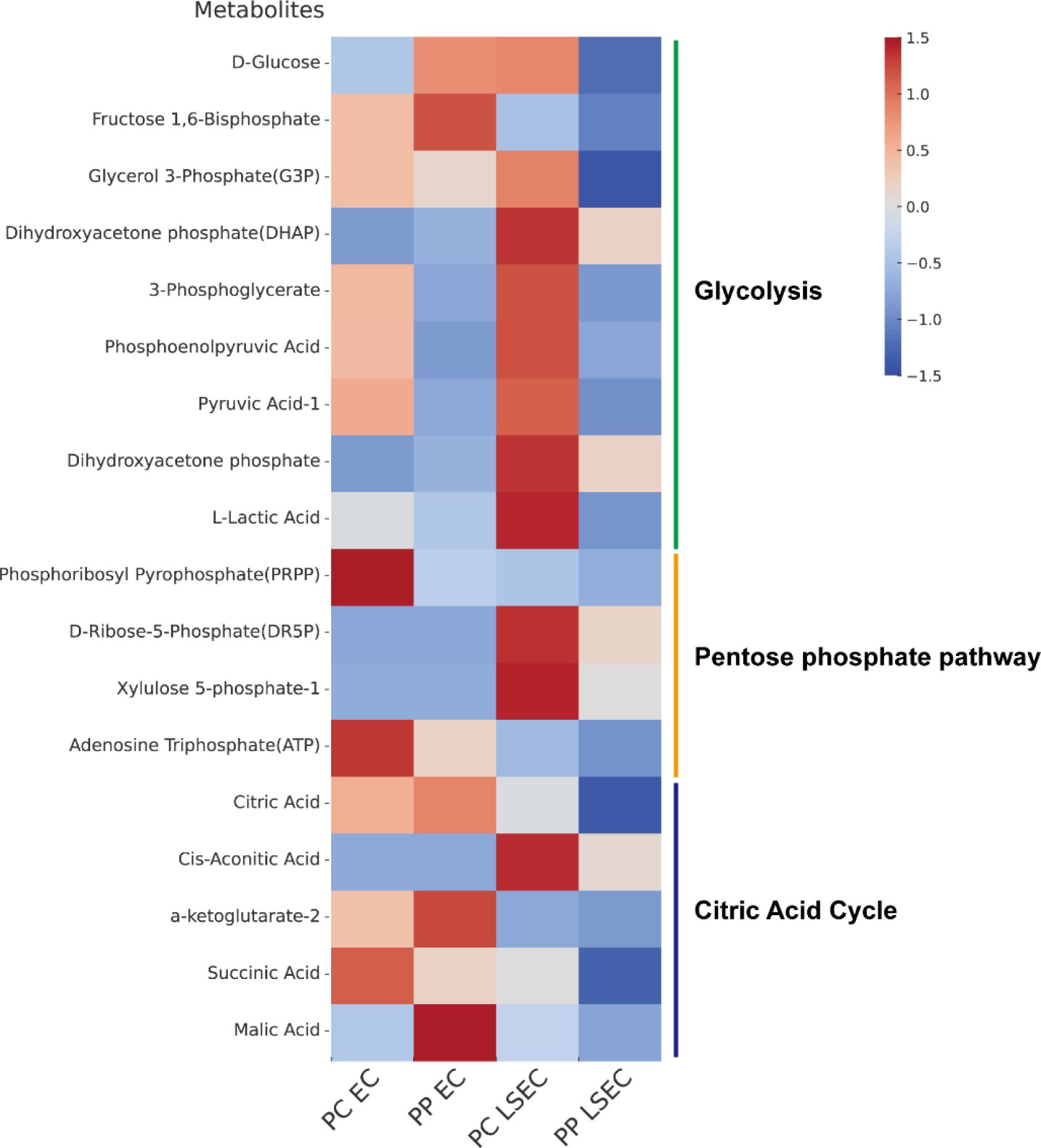
Metabolic profiling reveals differential glycolysis, pentose phosphate and tricarboxylic acid cycle metabolite activity in PP/PC EC/LSECs, related to Figure 1. Using mass spectrometry-based metabolomics, we analyzed the concentration of key glucose metabolites in PC ECs and PP LSECs compared to PP ECs and PP LSECs. PC EC and PC LSEC displayed significantly elevated levels of glycolytic and PPP intermediates, including glycerol-3-phosphate(G3P), fructose-6-phosphate (F6P), lactate, glucose-6-phosphate (G6P), phosphoribosyl pyrophosphate (PRPP), deoxyribose-5-phosphate (DR5P) and others, indicative of heightened glycolytic and PPP flux compared to PP EC and PP LSEC.

**Sup Figure 3.**
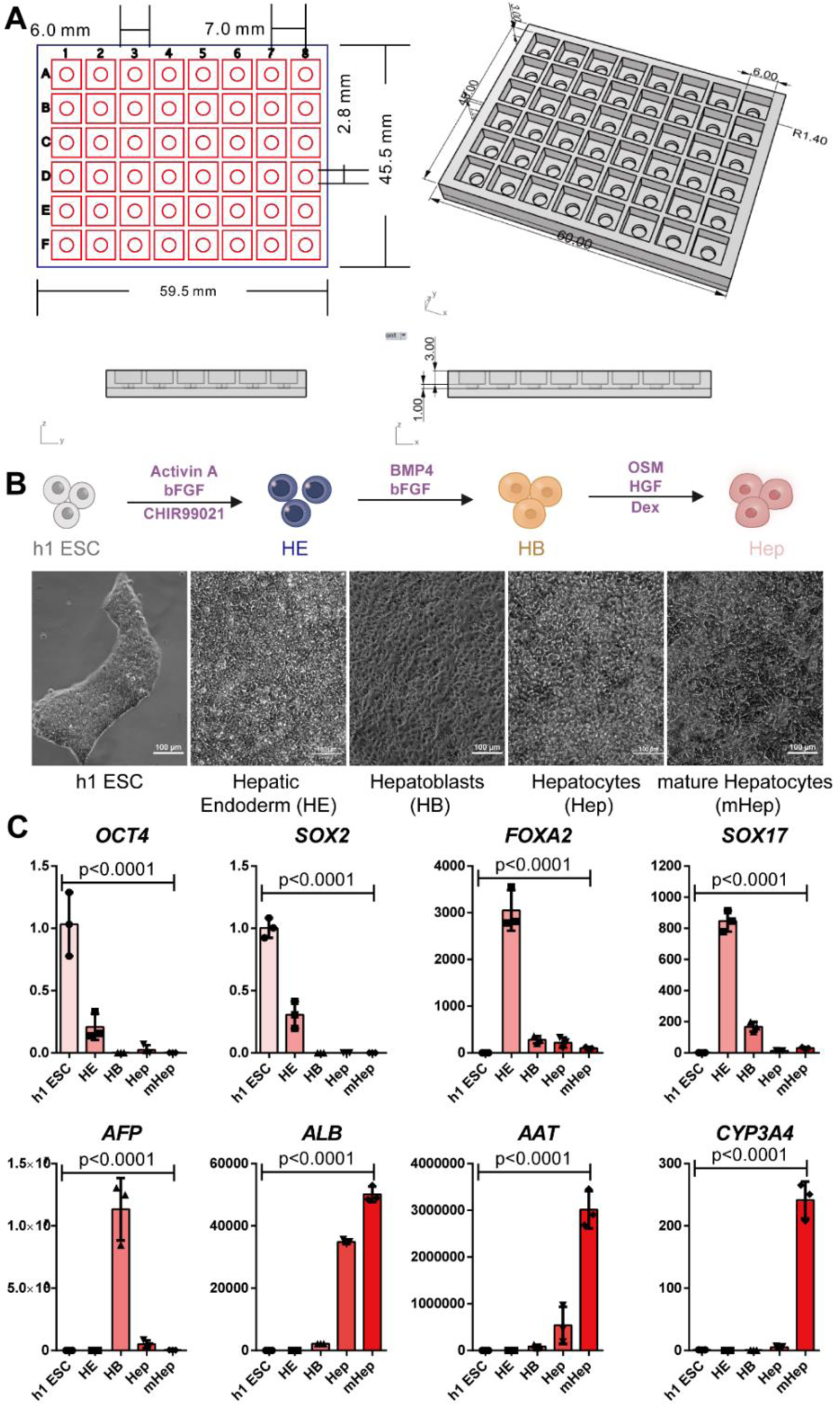
Microwell chip and cellular materials for 3D culture, related to Figure 2. (A) Schematic and three-dimensional diagrams of the cell culture plate, with specific dimensional parameters labeled on the illustrations; (B) The differentiation process of hepatocytes and the morphological changes at each stage; (C) qPCR Analysis of hepatocyte differentiation markers. OCT4 and SOX2 for hESC, FOXA2 and SOX17 for hepatic endoderm (HE), AFP for hepatoblasts (HB), ALB for hepatocytes (Hep), and AAT and CYP3A4 for more mature hepatocytes (mHep).

**Sup Figure 4.**
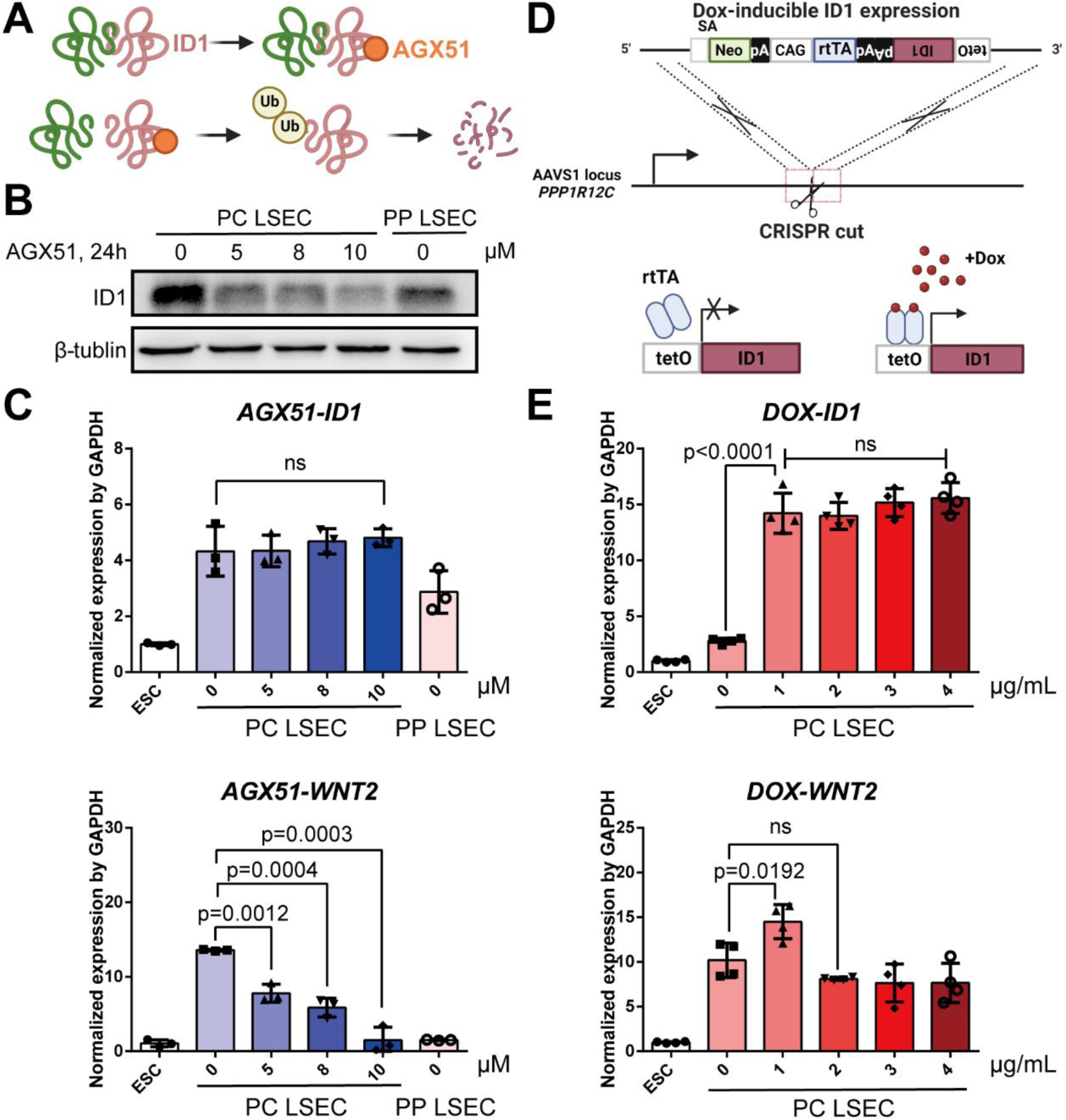
The correlation between ID1 and WNT2, related to Figure 4. (A) The schematic illustrates that AGX51 inhibitors promote the degradation of ID1 by facilitating the ubiquitination of the ID1 protein; (B) Western blot analysis for ID1 protein in hESC-derived PC and PP LSEC treated with different doses of AGX51, with β-tublin as a housekeeping protein; (C) qPCR results for the expression of *ID1* and *WNT2* genes in hESC-derived PC and PP LSEC treated with different doses of AGX51, with *GAPDH* as a housekeeping gene; (D) The schematic illustrated the construction of the Dox-inducible ID1-overexpressing Tet-on system in hESC cell line. (E) qPCR results for the expression of *ID1* and *WNT2* genes in ID1 KI hESC-derived PC LSEC treated with different doses of DOX, with *GAPDH* as a housekeeping gene.

**Sup Figure 5.**
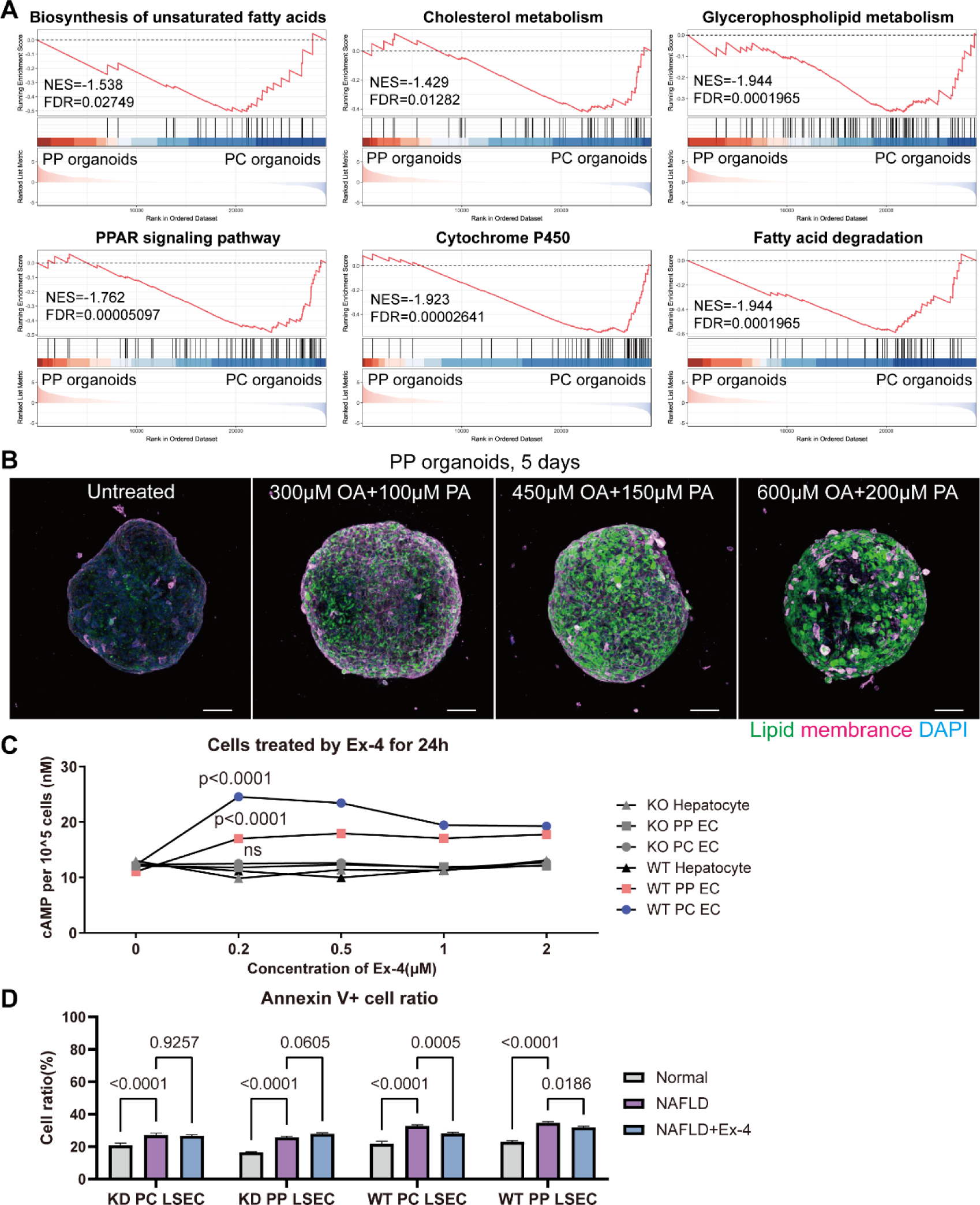
Results related to Figure 5. (A) Gene Set Enrichment Analysis (GSEA) of NAFLD and lipid metabolism pathways in PC and PP Organoids; (B) Characterization of lipid accumulation in PP organoids after treatment with different doses of FFAs for 5 days. Cell membrance is visualized in green, Lipid accumulation is visualized in green. The combination of 450 μM OA and 150 μM PA represents the most suitable conditions for inducing NAFLD; (C) Quantitative analysis of cAMP production in WT hESC-derived PC LSEC, PP LSEC, hepatocyte, and GLP-1R KD hESC-derived PC LSEC, PP LSEC, hepatocyte treated with various concentrations of Ex-4; (I) Flow cytometry quantitative analysis of apoptotic signal Annexin V in WT hESC-derived PC and PP LSEC, and GLP-1R KD hESC-derived PC and PP LSEC treated with Normal, NAFLD, NAFLD+ 200 nM Ex-4 medium for 24h.

**Sup Figure 6.**
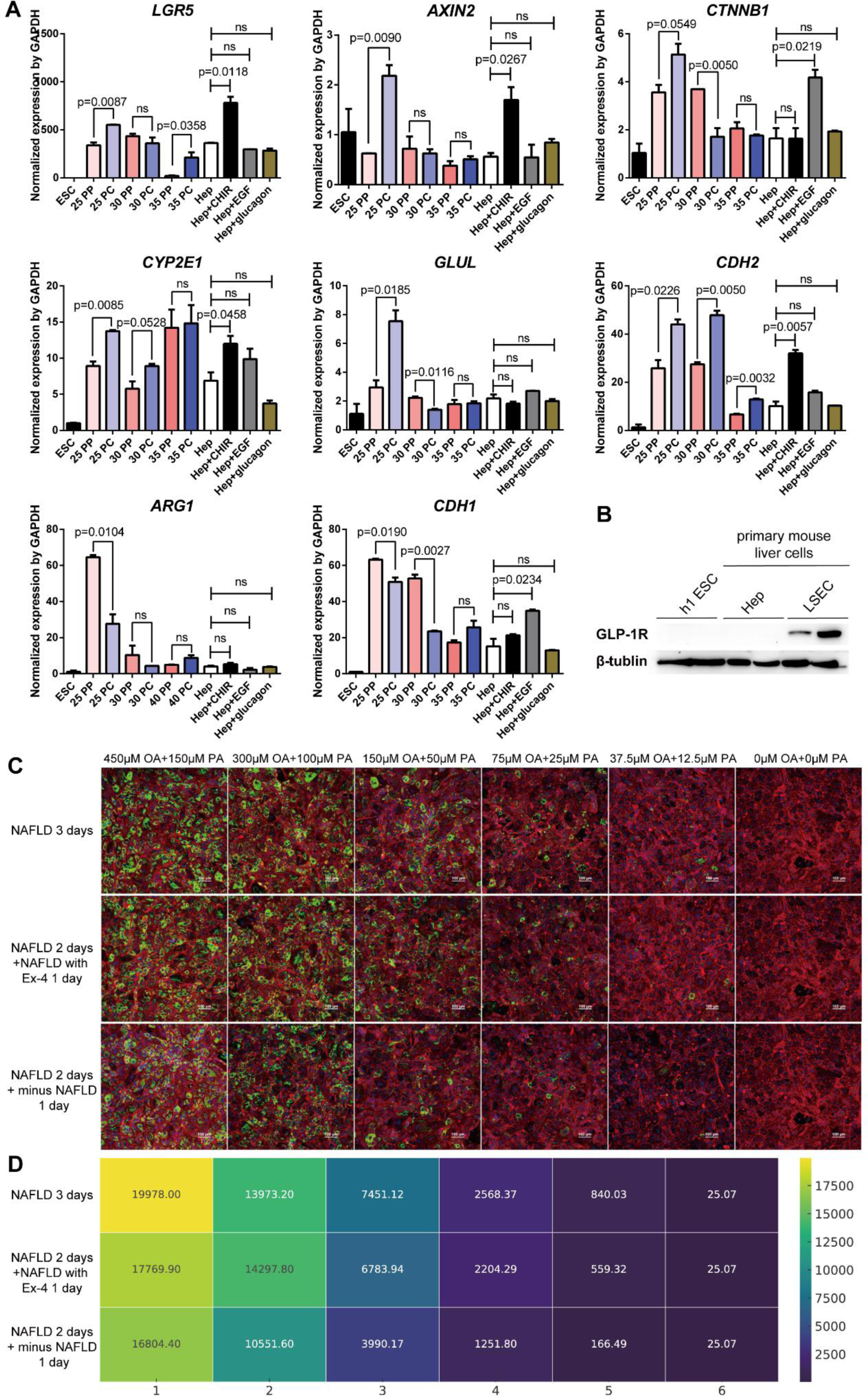
Results related to discussion. (A) qPCR results for the expression of downstream genes of the WNT/β-catenin signaling pathway (*LGR5*, *CTNNB1*, *AXIN2*), primary liver PC zone markers (*GLUL*, *CYP2E1* and *CDH2*) and primary liver PP zone markers (*ARG1* and *CDH1*) in PC and PP organoids, with *GAPDH* as a housekeeping gene. 3D organoid sample include 25PP (day25 hepatocyte co-culture with PP LSEC), 25PC (day25 hepatocyte co-culture with PC LSEC), 30PP (day30 primary mature hepatocyte co-culture with PP LSEC), 30PC (day30 primary mature hepatocyte co-culture with PC LSEC), 35PP (day35 mature hepatocyte co-culture with PP LSEC), 35PC (day35 mature hepatocyte co-culture with PC LSEC). 2D cell sample include hESC, hepatocyte, hepatocyte treated with 2 μM CHIR99021, 20 ng/mL EGF, 5 ng/mL glucagon. (B) Western blot analysis for GLP-1R protein in hESC, primary mouse LSEC and hepatocyte, with β-tublin as a housekeeping protein. (C) Characterization and quantification (D) of lipid accumulation in 2D hepatocyte co-culture with LSEC treated with various concentrations of FFAs and FFAs plus 200 nM Ex-4. Lipid accumulation is visualized in green, while F-actin, characterizing the cytoskeleton, is depicted in red. “Minus NAFLD” refers to the use of lower concentrations of FFAs in adjacent conditions.

**Table S1.**
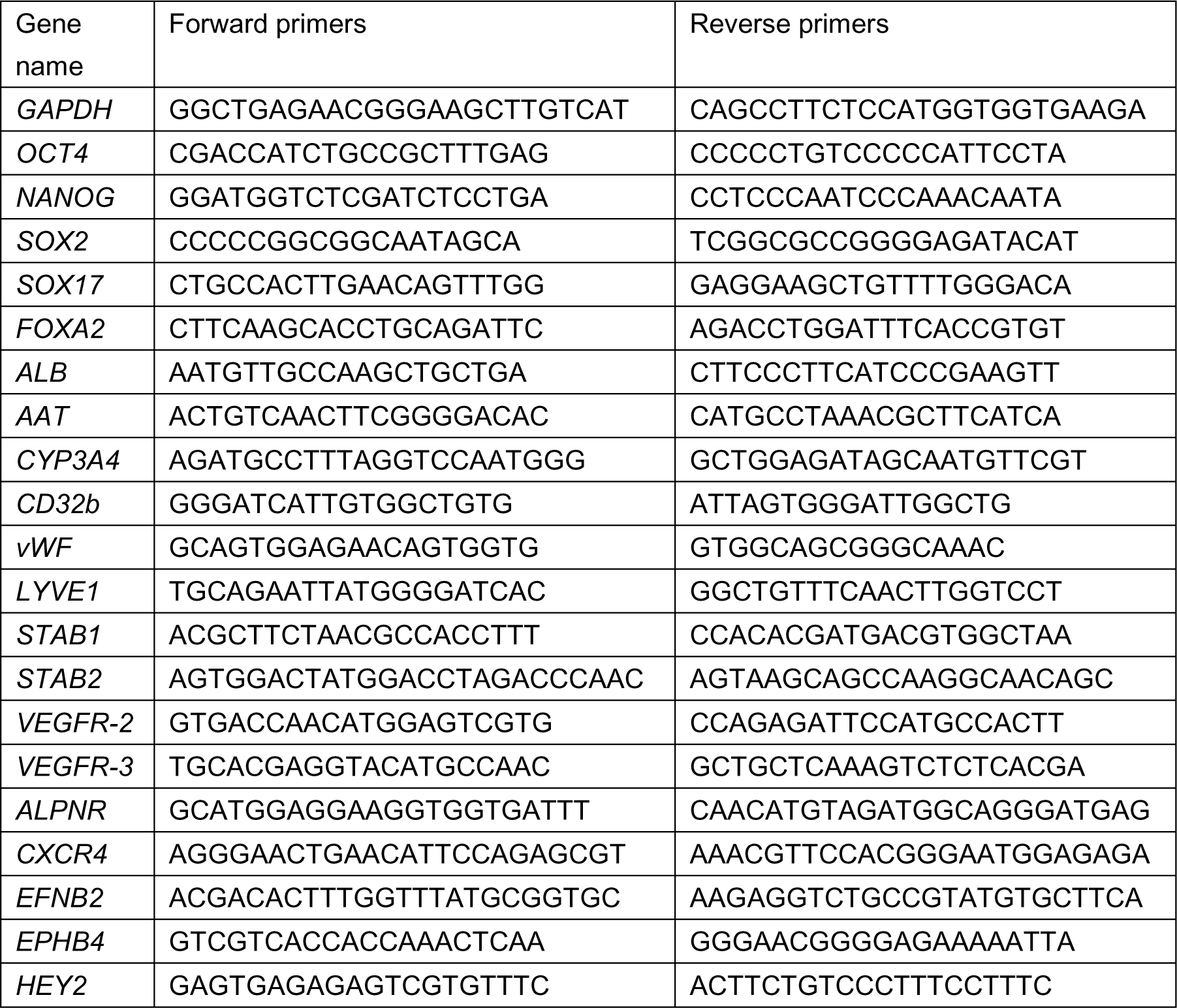
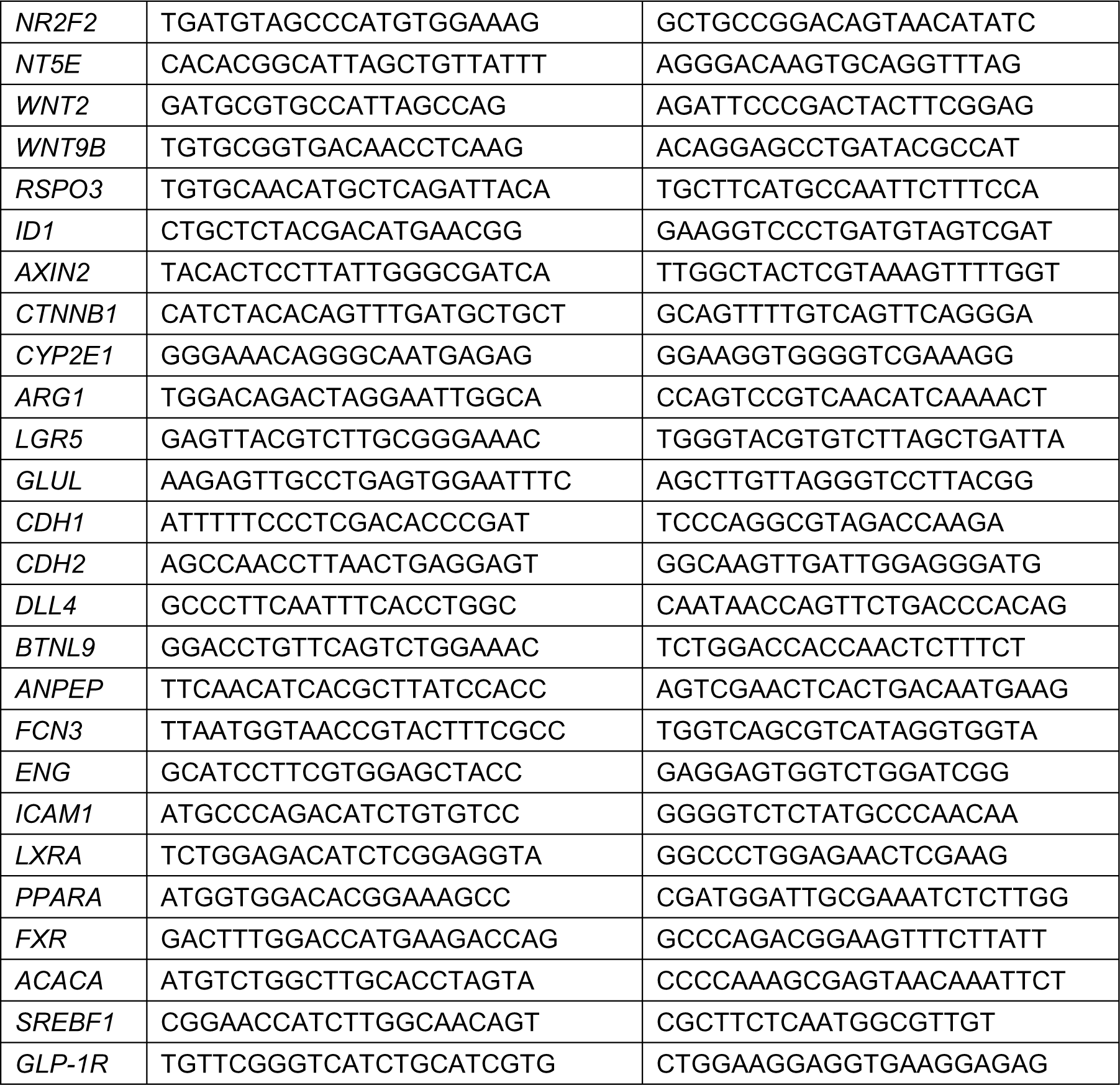
Primers for qPCR.

## Notes

### Competing Interest Statement

The authors have declared no competing interest.

